# Active synthesis of type I collagen homotrimer in Dupuytren’s fibrosis is unaffected by anti-TNF-α treatment

**DOI:** 10.1101/2020.07.13.195107

**Authors:** Kate Williamson, Katie J. Lee, Emma L. Beamish, Alan Carter, Jade A. Gumbs, Gabriella Cooper, Graham Cheung, Daniel Brown, Rob Pettitt, Eithne J. Comerford, Peter D. Clegg, Elizabeth G. Canty-Laird

## Abstract

Dupuytren’s disease is a common fibroproliferative disease of the palmar fascia of the hand with advanced cases treated surgically. Anti-tumour necrosis factor (TNF) injection has undergone phase 2 trials and may be effective in slowing early-stage disease progression. Here we sought to determine how new synthesis of type I collagen in Dupuytren’s differs from normal palmar fascia samples and to analyse the role of TNF in aberrant collagen synthesis. Model non-fibrotic, but fibrous connective tissues, were used to analyse active type I collagen protein synthesis in development, ageing and degenerative disease, where it was restricted to early development and ruptured tissue. Dupuytren’s tissue was shown to actively synthesise type I collagen, including abnormal type I collagen homotrimer. TNF-α reduced *COL1A2* gene expression only in the presence of serum in 2D cell culture and had opposing effects on collagen protein production in the presence or absence of serum. TNF-α had only limited effects in 3D tendon-like constructs. Anti-TNF did not reduce type I collagen synthesis in 3D tendon-like constructs or prevent type I collagen homotrimer synthesis in Dupuytren’s tissue. Hence, modulation of the TNF-α pathway in Dupuytren’s disease is unlikely to prevent the pathological collagen accumulation that is characteristic of fibrosis.

## Introduction

Fibrillar collagens, particularly type I, are the major structural component of fibrous connective tissue including palmar fascia (aponeurosis), tendon, ligament, and fibrotic tissue. In fibrous tissue string-like fibrils comprise arrays of collagen molecules. Excessive accumulation of fibrillar collagens impedes normal tissue function resulting in particularly poor outcomes in cardiac, pulmonary, kidney and liver fibrosis (Wynn, 2008; Zeisberg and Kalluri, 2013). Dupuytren’s disease is a common fibroproliferative disorder of the palmar fascia of the hand, which has sex, age, geographic and racial differences; being most prevalent in, but not restricted to, older men of Northern European descent (Hindocha et al., 2009). The aetiology of Dupuytren’s is complex involving genetics; an autosomal dominance pattern with varying penetrance, links to other diseases such as diabetes, epilepsy and liver disease, and environmental factors including alcohol intake and smoking (Grazina et al., 2019).

The formation of fibrous tissue under the skin can cause Dupuytren’s patients localised pain and discomfort whilst disease progression may prevent the digits from straightening, producing fixed flexion contractures. The disease severity is commonly measured by the degree of joint contraction (which can be monitored by Tubiana staging), the number of digits involved and the presence of disease outside the hand; Ledderhose disease in the foot or Peyronie’s disease in the penis (Dibenedetti et al., 2011; Hindocha et al., 2008). Treatment options include surgery; fasciectomy (excising the diseased tissue) and dermofasciectomy (excising the disease tissue and overlying skin), percutaneous needle aponeurotomy (dividing the diseased tissue) and collagenase treatment (prior to withdrawal from several jurisdictions) (Soreide et al., 2018). Each treatment results in varying success, complications and recurrence rates (Krefter et al., 2017). In patients with advanced disease and debilitating contractures, the gold standard treatment is open limited fasciectomy followed by physiotherapy to encourage improved range of movement (Worrell, 2012). Surgical complication rates increase with disease severity and range between 3-50% (Craxford and Russell, 2016), whilst recurrence in the same finger/thumb or disease progression in other digits is common (8-54%) (Worrell, 2012).

The fibrotic tissue in Dupuytren’s can comprise a highly cellular ‘nodule’ region, which is thought to represent an active stage in the tissue pathogenesis and a ‘cord’ region, consisting of mature fibrillar collagen (Layton and Nanchahal, 2019). Disease pathogenesis has been divided into proliferative, involutional (contracting) and residual stages, with progressively decreasing cellularity and increasing alignment along directional lines of tension (Luck, 1959). Type III collagen is present at very low levels in normal palmar fascia, but is abundant in Dupuytren’s tissue (Brickley-Parsons et al., 1981). However, the proportion of type III collagen relative to total collagen decreases through the stages of disease progression from >35% to <20% in the residual stage (Lam et al., 2010). Myofibroblasts are abundant in nodules, display persistent alpha-smooth muscle actin (alpha-SMA) expression and are responsible for both matrix deposition and contraction. Bidirectional actin-fibronectin interactions (Tomasek and Haaksma, 1991) result in the progressive tensioning of collagen fibres and a concurrent increase in total flexion deformity (Verjee et al., 2009). Other cell types implicated in the initiation or progression of Dupuytren’s fibrosis include embryonic stem cells, mesenchymal stromal cells, fibrocytes and immune cell populations (Layton and Nanchahal, 2019; Tan et al., 2018).

Analysis of messenger RNA (mRNA) expression profiles in isolated Duputyren’s fibroblasts found increased collagen and extracellular matrix (ECM) mRNAs (Forrester et al., 2013), whilst a loss of collagen-regulating microRNAs (miRs) was identified in Dupuytren’s tissue (Riester et al., 2015). A weighted gene co-expression network analysis and functional enrichment analysis of Dupuytren’s transcriptomic datasets found gene ontology terms for ECM and collagen in ECM organisation, ECM-receptor interaction and collagen catabolic process (Jung et al., 2019). Type I collagen is the major ECM component of fibrotic Dupuytren’s tissue and type I collagen molecules are predominantly (α1)_2_(α2)_1_ heterotrimers derived from the polypeptide gene products of the *COL1A1* and *COL1A2* genes. However, abnormal collagen (α1)_3,_ homotrimer has been identified in Dupuytren’s tissue (Ehrlich et al., 1982). Type I collagen homotrimers are resistant to proteolytic degradation by matrix metalloproteinases (MMPs) (Makareeva et al., 2010) and may therefore skew the balance between type I collagen synthesis and degradation, impeding the resolution of fibrosis.

A systematic review of Dupuytren’s disease ‘omics’ studies highlighted alterations in collagen and ECM gene expression as well as genomic and transcriptomic studies implicating the TGF-β and Wnt signalling pathways in disease pathogenesis (Shih et al., 2012). TNF at 0.1 ng/ml was found to induce cell contractility, to increase a-SMA mRNA and protein, and to increase COL1A1 mRNA in palmar fibroblasts, from Dupuytren’s patients; effects that could be reversed with the Wnt pathway inhibitor lithium chloride at 10mM (Verjee et al., 2013). Conversely, anti-TNF reduced contractility, led to disassembly of the actin cytoskeleton, reduced a-SMA mRNA and protein, and decreased COL1A1 mRNA in Dupuytren’s myofibroblasts derived from nodular tissue. Subsequently anti-TNF therapy (adalimumab) in Dupuytren’s patients delivered by intra-nodular injection of a 40 mg dose was found to reduce a-SMA and type I procollagen protein production (as an N-propeptide assay), though not *COL1A1* gene expression, or total collagen, in nodules obtained during surgery 2 weeks after injection (Nanchahal et al., 2018). A longer-term study comprising four injections every three months showed a reduction in nodule size and nodule softening (Nanchahal et al., 2022) in patients with early-stage Dupuytren’s disease. Prior studies have shown that TNF-α decreases, rather than increases *COL1A1* gene expression and collagen protein synthesis in human dermal fibroblasts at 0.1ng/ml and higher concentrations (Mori et al., 1996; Takeda et al., 1993). COL1A2 expression and type I collagen protein is similarly decreased with 10 ng/ml TNF-α in immortalised mouse fibroblasts, COL1A2 expression decreased in human dermal fibroblasts (Verrecchia et al., 2002) and Col1a1 expression decreased in rat hepatic stellate cells (Varela-Rey et al., 2002).

We hypothesised that there may be a continual production of abnormal homotrimeric type I collagen that could contribute to the recurrence of Dupuytren’s contracture following medical treatment. Whilst TNF has been identified as a potential therapeutic target in Dupuytren’s disease (Verjee et al., 2013), information on the response of Dupuytren’s cells and tissues to TNF-α in terms of collagen protein synthesis and production of normal versus abnormal type I collagen is currently lacking. This information is critically important given that type I collagen protein is by definition the major constituent of fibrotic tissue.

In this study, we utilised metabolic labelling approaches in combination with 1D gel electrophoresis to analyse type I collagen synthesis in fibrotic Dupuytren’s tissue, and in non-fibrotic, fibrous connective tissue. The study aimed to determine how fibrotic collagen differs from that produced in normal fibrous tissue and how type I collagen synthesis differs from that seen in development, or in injured tissue. We evaluated the type I collagen biosynthetic response of normal palmar fascia fibroblasts, and those from Dupuytren’s nodule and cord to TNF-α in 2D cell culture, and in cell-assembled collagenous 3D cultures. The effect of anti-TNF treatment on type I collagen biosynthesis was also evaluated in tissue explants and 3D cultures.

## Results

### Type I collagen homotrimer is actively produced by Dupuytren’s tissue

To determine if type I collagen homotrimer is continuously synthesized in Dupuytren’s tissue, *COL1A1* and *COL1A2* gene expression was measured by qRT-PCR, and synthesis of α-chains by radiolabelling, in Dupuytren’s samples (Supplementary Table 1) and in normal palmar fascia (PF) controls (Supplementary Table 2). Expression of COL1A1 (Fig. 1A) and COL1A2 (Fig. 1B) were significantly higher in Dupuytren’s tissue (p = 0.001 and 0.002 respectively), indicative of new type I collagen synthesis. The relative expression of COL1A1 as compared to COL1A2 was also higher in Dupuytren’s tissue (p = 0.048) (Fig. 1C) indicating increased gene transcription and/or mRNA stability of COL1A1 as compared to COL1A2. There was no significant difference in the age of the Dupuytren’s and normal PF samples that were analysed by qRT-PCR (p=0.11).

**Figure 1:**
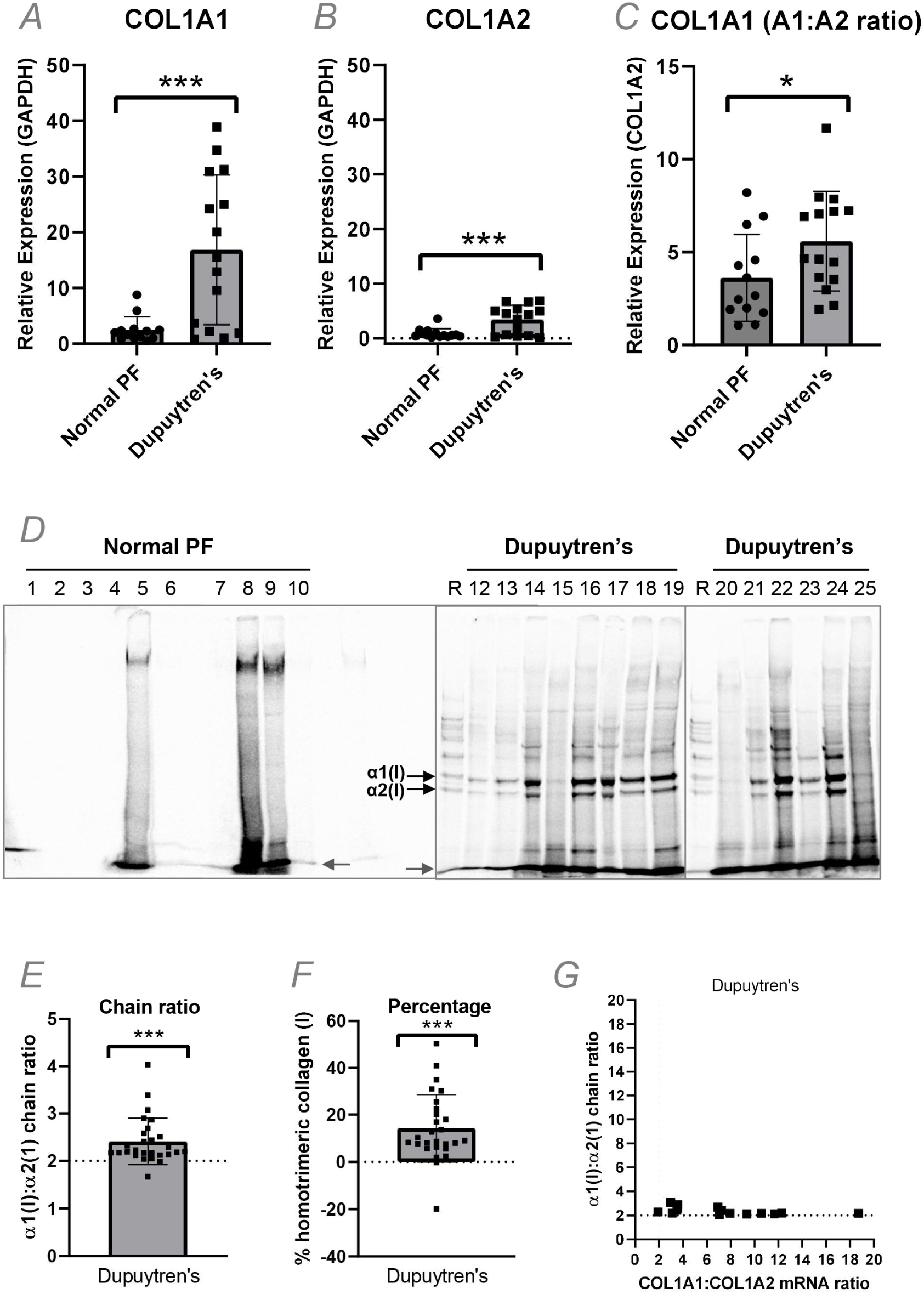
Synthesis of type I collagen homotrimer by Dupuytren’s tissue. A-C: Analysis of *COL1A1* (A), *COL1A2* (B) and COL1A1:COL1A2 (C) gene expression by qRT-PCR in normal PF (n=13) and Dupuytren’s tissue samples (n=15). D: Representative delayed reduction 6% SDS-Page gels of normal PF (D) and Dupuytren’s (E) tissue extracts after pulse-chase labelling with [^14^C]proline. E-F: The relative amounts of the labelled α1(I) and α2(I) chains in Dupuytren’s (n=27) samples were quantified by densitometry and expressed as an α1(I):α2(I) chain ratio (E) or converted to a percentage of homotrimeric type I type I collagen (F). The α1(I):α2(I) chain ratio was plotted against the COL1A1:COL1A2 mRNA ratio for Dupuytren’s samples (n=12) for which both data types were available (G). * p<0.05, ** p<0.01 and *** p<0.001.

Metabolic labelling with ^14^C proline indicated Dupuytren’s tissue explants but not normal PF controls (n=22) produced newly-synthesised type I collagen protein (Fig. 1D). Samples derived from 27 of 31 Dupuytren’s patients were sufficiently labelled for densitometric quantification (Supplementary Table 1). The α1(I):α2(I) chain ratio was significantly greater than 2 (p<0.001) (Fig. 1E) indicative of new type I collagen homotrimer synthesis. Using the ratios for the nodule only in the analyses (discounting 3 other cord samples) produced similar results (p<0.001, data not shown). The α1(I):α2(I) ratio was converted to a percentage of homotrimeric type I collagen, indicating a mean value of 14.3% (SD ± 14.4%) and with one sample reading 50.4% (Fig. 1F). Plotting the α1(I):α2(I) ratio against the COL1A1:COL1A2 mRNA ratio indicated no positive linear relationship between relative mRNA and polypeptide chain ratios (Fig. 1G), and the Pearson correlation coefficient was negative (r = -0.522, p = 0.046).

### Demographic factors are not associated with greater type I collagen homotrimer synthesis

To determine if particular demographic factors were associated with an increased proportion of homotrimeric type I collagen, a Principal Components Analysis (PCA) was carried out (Fig. 2). Samples producing lower (<10%), medium (10-25%) or higher (>25%) percentages of homotrimeric type I collagen did not group based on demographics factors (Fig. 2A). Demographic factors (Fig. 2B) had a minimal effect on principal component scores. To determine whether disease stage influences type I collagen homotrimer synthesis, the PCA was repeated on samples with paired disease stage, which also showed only a weak effect (Fig. 2C-D).

**Figure 2:**
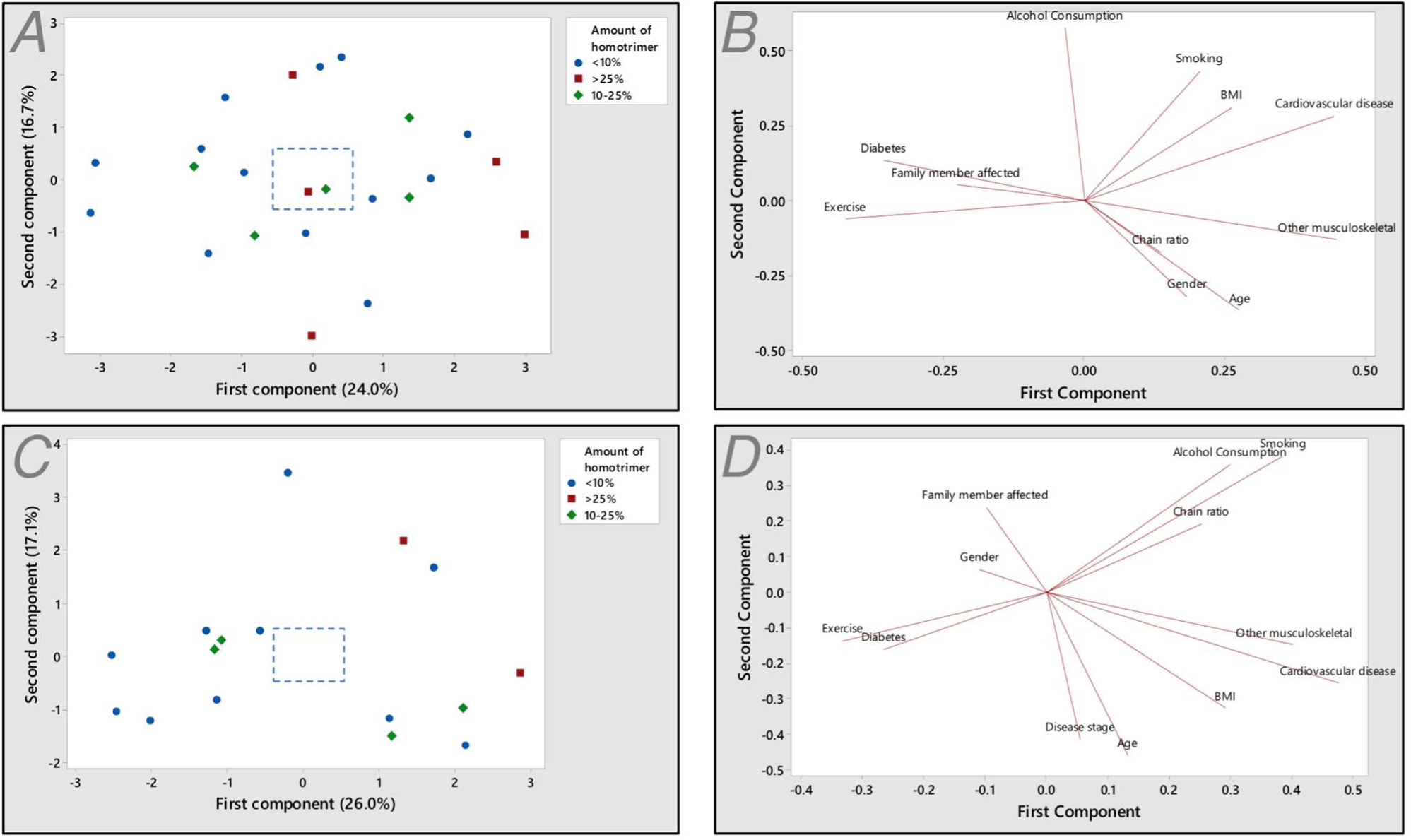
Principal components analysis of the relationship between demographic factors and the proportion of homotrimeric collagen synthesized by human Dupuytren’s surgical samples. A: Score plot grouped by lower (<10%), medium (10-25%) and higher (>25%) percentages of type I collagen homotrimer (n=23). The box indicates the relative scale on the loading plot (B). C-D: Score plot (C) and loading plot (D) for a patient sub-set with disease stage information (n=16).

### Active type I collagen synthesis in diseased, ruptured or foetal, but not healthy adult fibrous connective tissue

No radiolabeled normal PF samples (Supplementary Table 2) were found to synthesise detectable amounts of type I collagen in explant culture, but a high molecular weight band was noted in 10 of the 22 labelled normal PF samples (examples shown in Fig. 1D). Under reducing conditions, a single band migrating similarly to the α2(I) chain was noted in normal PF samples and one between the α1(I) and α2(I) chain in some Dupuytren’s samples (#s 12, 14 and 19) (Fig. 3A). The identity of the labelled proteins in these bands are unknown but are expected to be collagenous given the incorporation of labelled proline.

**Figure 3:**
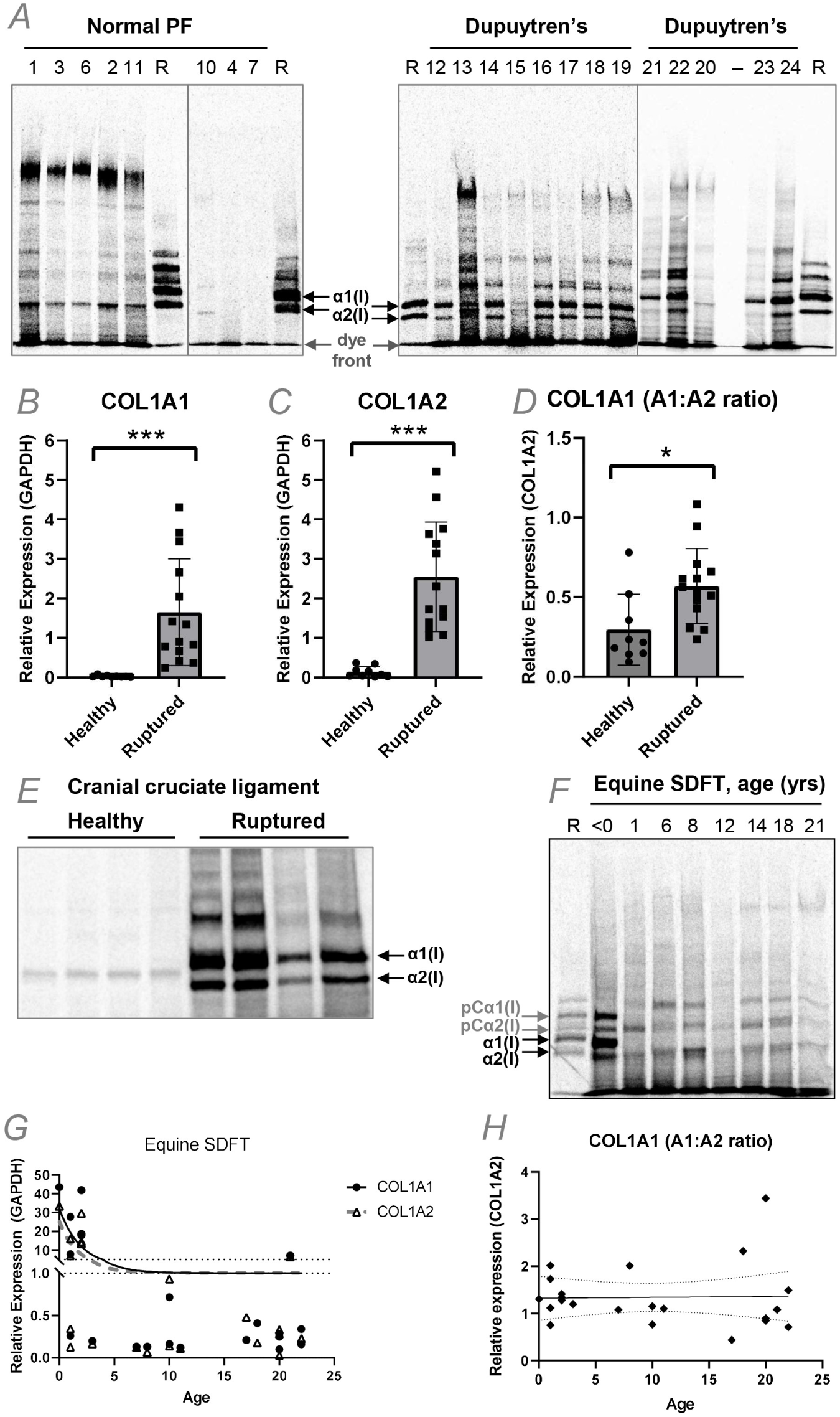
Type I collagen and proline-rich protein synthesis in tendon and ligament explants. A: Representative 4% SDS-PAGE gels of reduced normal PF and Dupuytren’s tissue extracts after pulse-chase labelling with [^14^C]proline. Sample numbers relate to those shown in Figure 1D. B-D: Analysis of *COL1A1* (B), *COL1A2* (C) and COL1A1:COL1A2 (D) gene expression by qRT-PCR in healthy (n=9) and ruptured (n=14) canine cranial cruciate ligament samples. E: Representative 4% SDS-Page gel of reduced healthy and ruptured canine cranial cruciate ligament extracts after pulse-chase labelling with [^14^C]proline. F: Representative 4% SDS-PAGE gel of equine superficial digital flexor tendon (SDFT) extracts at different ages after pulse-chase labelling with [^14^C]proline. G-H: Analysis of *COL1A1* and *COL1A2* (G) and COL1A1:COL1A2 (H) gene expression by qRT-PCR in equine superficial digital flexor tendon (SDFT) at various ages. Trend lines and 95% confidence intervals (H) are shown. Note; horses have an average 25-30 year lifespan, with 20 years being equivalent to approximately 60 human years.

We considered that the age and normal status of the normal PF samples may preclude detectable levels of new type I collagen synthesis. Model non-fibrotic fibrous connective tissues were therefore studied. Analysis of canine cranial cruciate ligament (CCL) allowed a comparison between healthy fibrous tissue, and those that were ruptured due to degenerative disease, whilst analysis of equine superficial digital flexor tendon (SDFT) facilitated a comparison across ages: foetal, young (yearling), adult (6-14 yrs) and old (>15 yrs). The relative expression of both COL1A1 (Fig. 3B) and COL1A2 (Fig. 3C) was significantly increased (both p<0.001) in ruptured canine cranial cruciate ligament as compared to healthy samples. The expression of COL1A1 relative to COL1A2 was higher in ruptured ligament (p = 0.012) (Fig. 3D), although the ratio did not exceed 1.5. Ruptured ligament produced newly-synthesised type I collagen whilst under reducing conditions healthy ligament produced a single band migrating between the α1(I) and the α2(I) chain (Fig. 3E). Foetal SDFT produced nascent type I collagen as expected, whilst all post-natal SDFT samples did not; instead producing a band co-migrating with α2(I) (similar to human normal PF), a higher molecular weight band co-migrating with pCα2(I) and often a band migrating between proα1(I) and pCα1(I) (Fig. 3F). Gene expression of *COL1A1* and *COL1A2* in post-natal samples was highest before 2 years of age (Fig. 3E) but there was no significant correlation between the COL1A1:COL1A2 gene expression ratio and age (Fig. 3H).

### TNF-α reduces *COL1A2* gene expression in the presence of serum in cell culture and has opposing effects on collagen protein production in the presence or absence of serum

To determine whether TNF-α increased or inhibited type I collagen synthesis in Dupuytren’s cells, cells derived from Dupuytren’s nodule, cord or from normal PF (Supplementary Table 3) were treated with 10 ng/ml TNF-α in the presence or absence of 10% serum. Without serum, COL1A1 and COL1A2 gene expression was significantly higher in nodule cells than normal PF (p<0.001 for both) and cord cells (p=0.004 and 0.008 respectively), but unaltered by TNF-α treatment (Fig. 4 A&B). The COL1A1:COL1A2 gene expression ratio was higher in cells from nodule than normal palmar fascia (p=0.026), but the ratio was consistently less than 2 and unaffected by TNF-α treatment (Fig. 4C).

**Figure 4:**
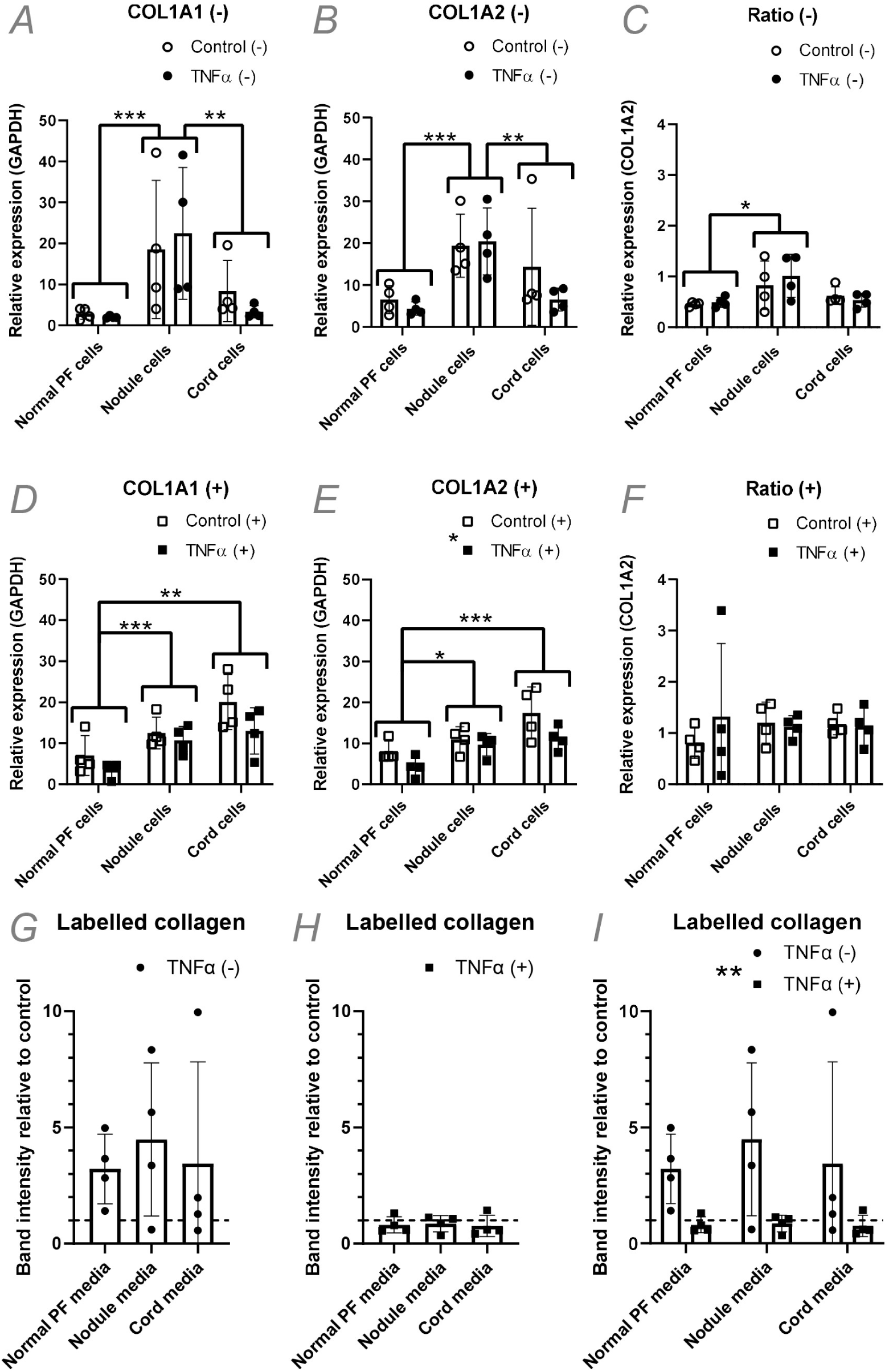
Effect of TNF-α on type I collagen gene expression and collagen protein production in Dupuytren’s cells. A-F: Analysis of *COL1A1* (A, D), *COL1A2* (B, D) and COL1A1:COL1A2 (C, F) gene expression by qRT-PCR in normal PF, Dupuytren’s nodule and Dupuytren’s cord cells (n=4) after TNF-α treatment in serum free conditions (-) (A-C) or in the presence of 10% FCS (+) (D-F). G-I: Densitometric quantification of the relative amounts of radiolabelled (pro)collagen present in conditioned media from normal PF, Dupuytren’s nodule and Dupuytren’s cord cells (n=4) after TNF-α treatment, as compared to control treatments, in serum free conditions (-) (G), the presence of 10% FCS (+) (H) or in both conditions (I). * p<0.05, ** p<0.01 and *** p<0.001. Sample details are given in Supplementary Table 3.

In the presence of serum, *COL1A1* and *COL1A2* gene expression was higher in both nodule (p=0.005 and p=0.015) and cord (p<0.001 for both) cells than in cells from normal palmar fascia (Fig. 4 D&E). TNF-α treatment significantly lowered *COL1A2* gene expression (p=0.019), but *COL1A1* gene expression was not significantly decreased (p=0.058). The COL1A1:COL1A2 gene expression ratios were unaffected (Fig. 4F).

There were no significant changes in the amount of radiolabeled (pro)collagen in the cell culture media following TNF-α treatment in either the absence (Fig. 4G) or presence (Fig. 4H) of serum as compared to control cultures, or between cell types. However, the mean values in TNF-α-treated cultures as compared to controls were all elevated in the absence of serum and decreased in the presence of serum. Comparing the effect of TNF-α treatment in the absence or presence of serum on the relative amount of labelled collagen in cell culture media showed a significant difference between conditions (p=0.001) (Fig. 4I).

### Normal PF and Dupuytren’s cells can form 3D tendon-like constructs that fully process type I collagen and assemble aligned collagen fibrils

Cells in 2D culture do not assemble a robust collagenous ECM (Canty and Kadler, 2005), although they do express type I collagen mRNA (Fig. 4A-F) and synthesize type I collagen trimers. The trimeric type I collagen is primarily produced in the procollagen form and secreted into the cell culture medium where it is partially processed (Fig. 4A). The cell layer is often poorly labelled with (pro)collagen being weakly distinguishable above background (not shown).

A fibrin-based 3D culture system was hence utilized in which cells replace the fibrin scaffold with a cell-derived collagenous ECM comprising bona-fide collagen fibrils and resembling embryonic tendon (Kapacee et al., 2010). Human tendon fibroblasts and MSCs have been previously shown to form such structures (Bayer et al., 2010). In the present study normal PF, Dupuytren’s nodule and cord cells (Supplementary Table 3) were grown in 3D culture and found to produce tendon-like constructs in 4 weeks. When labelled these constructs were found to produce fully processed type I collagen (Fig. 5B), whilst labelled procollagen and processing intermediates were present in the cell culture media (Fig. 5C). Ultrastructural analysis of the tendon-like constructs was performed using transmission electron microscopy (TEM) (Fig. 5 D-O & S1 A-F). Semi-thin sections showed the formation of dense cellular structures (Fig. 5 D-F & S1 A-C); similar in appearance to tissues formed from fibroblasts and MSCs (Bayer et al., 2014; Janvier et al., 2020; Kapacee et al., 2008). Transmission electron micrographs showed the presence of narrow diameter aligned collagen fibrils between cells (Fig 5 G-I & S1 D-F) also consistent with previous reports (Bayer et al., 2014; Janvier et al., 2020; Kapacee et al., 2008; Kharaz et al., 2016). Tissues derived from nodular fibroblasts appeared more compact with well-defined fibrils (Fig 5 E&H, S1 B&E).

**Figure 5:**
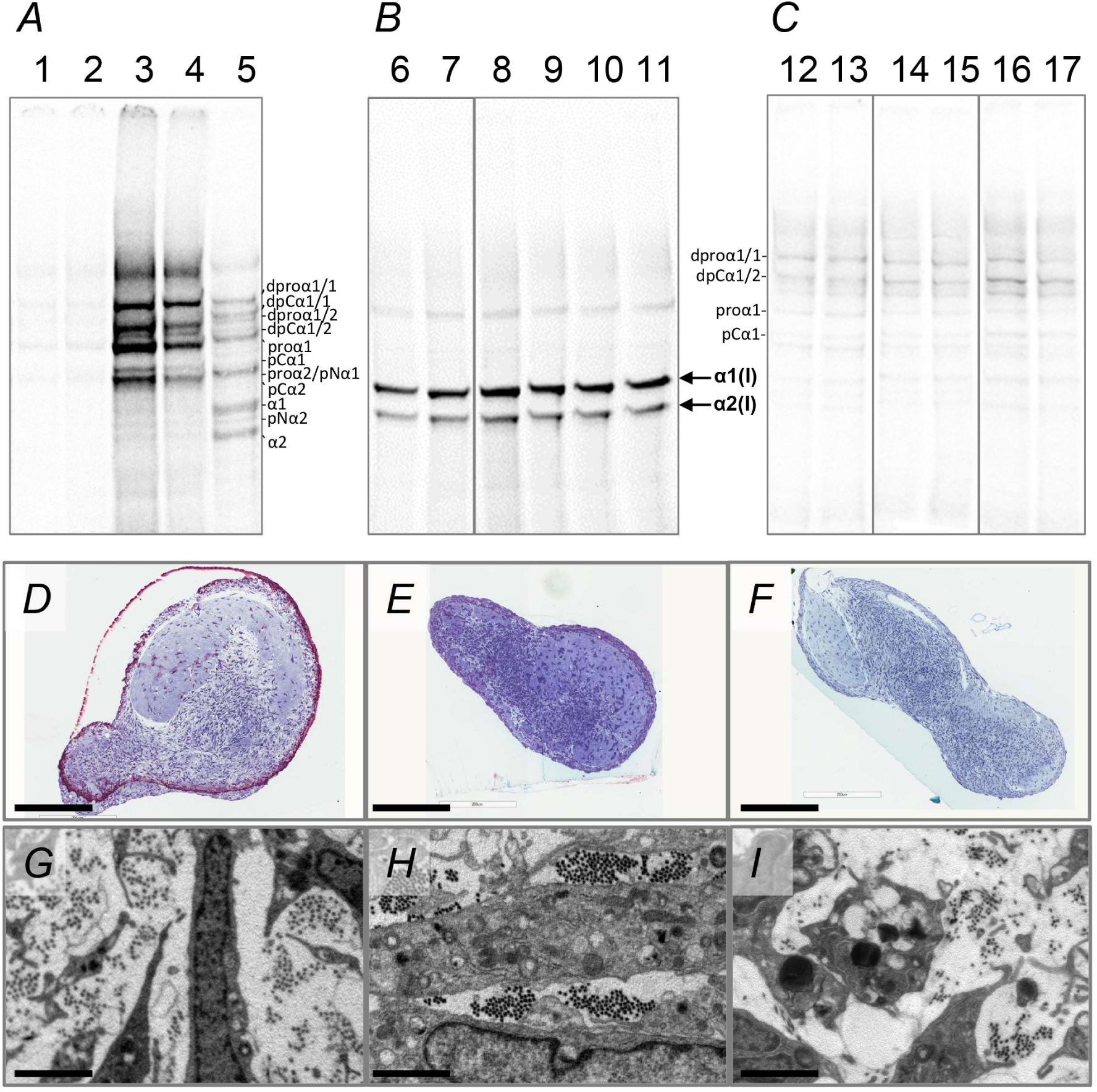
Normal palmar fascia and Dupuytren’s cells fully process procollagen, assemble *de novo* collagen fibrils and create tendon-like structures in 3D culture. A-C: Representative gel images of delayed reduction electrophoresis of the media (A&C) or salt extractions (B) of normal palmar fascia and Dupuytren’s cells grown in 2D (A) or 3D (B&C) culture and labelled with [^14^C]proline. Fully processed collagen was found in extracellular extracts of the tendon-like constructs (B), whereas pro-forms were found in the media of both 2D (A) and 3D cultures (C). The type I procollagen processing intermediates were identified by comparison to labelled standards analysed under reducing, non-reducing and delayed reduction conditions and are indicated; dproα1/1 (disulphide linked dimer of two proα1 chains), dpCα1/1 (disulphide linked dimer of two pCα1 chains), dproα1/2 (disulphide linked dimer of one proα1 and one proα2 chain), dpCα1/2 (disulphide linked dimer of one pCα1 and one pCα2 chain), proα1 (procollagen α1 chain), pCα1 (pC collagen α1 chain, missing the N-propeptide), proα2/pNα1 (co-migrating procollagen α2 chain/pN collagen α1 chain, missing the C-propeptide), pCα2 (pC collagen α2 chain, missing the N-propeptide), α1 (fully processed α1 chain), pNα2 (pN collagen α2 chain, missing the C-propeptide), α2 (fully processed α2 chain). For A-C lanes 1, 8 and 14 are the control for lanes 2, 9 and 15 which were treated with TNF-α in the absence of serum, and lanes 3, 10 and 16 are the control for lanes 4, 11 and 17 which were treated with TNF-α in the presence of serum. Samples in lanes 6 and 12 were treated with IgG control and in lanes 7 and 13 with anti-TNF-α. Lane 5 is a labelled type I (pro)collagen standard. D-F: Semi-thin toluidine blue-stained sections of 3D tendon-like constructs (bar 200 um). G-I: Transmission electron microscopy images of 3D tendon-like constructs (bar 2 um). Constructs were derived from normal palmar fascia cells (D&G), Dupuytren’s nodule (E&H) or Dupuytren’s cord (F&I). Sample numbers are given in Supplementary Table 3.

### TNF-α has limited effects on type I collagen gene expression and does not affect protein production in 3D tendon-like constructs

Tendon-like constructs derived from normal PF, Dupuytren’s nodule or cord cells (Supplementary Table 3) were treated with 10 ng/ml TNF-α in the presence or absence of 10% serum and type I collagen gene expression monitored by qRT-PCR (Fig 6 A-F). TNF-α had no effect on *COL1A1* or *COL1A2* gene expression in the presence or absence of serum (Fig 6 A-B & C-D), though *COL1A1* gene expression was lower in 3D constructs derived from normal PF cells than those derived from Dupuytren’s nodule or cord cells (p=0.033 and 0.039 respectively) (Fig 6A). There was no significant difference in the COL1A1:COL1A2 gene expression ratio with TNF-α treatment in the absence of serum (Fig. 6C), but the ratio was decreased in the presence of 10% serum (p=0.048) (Fig. 6F).

**Figure 6.**
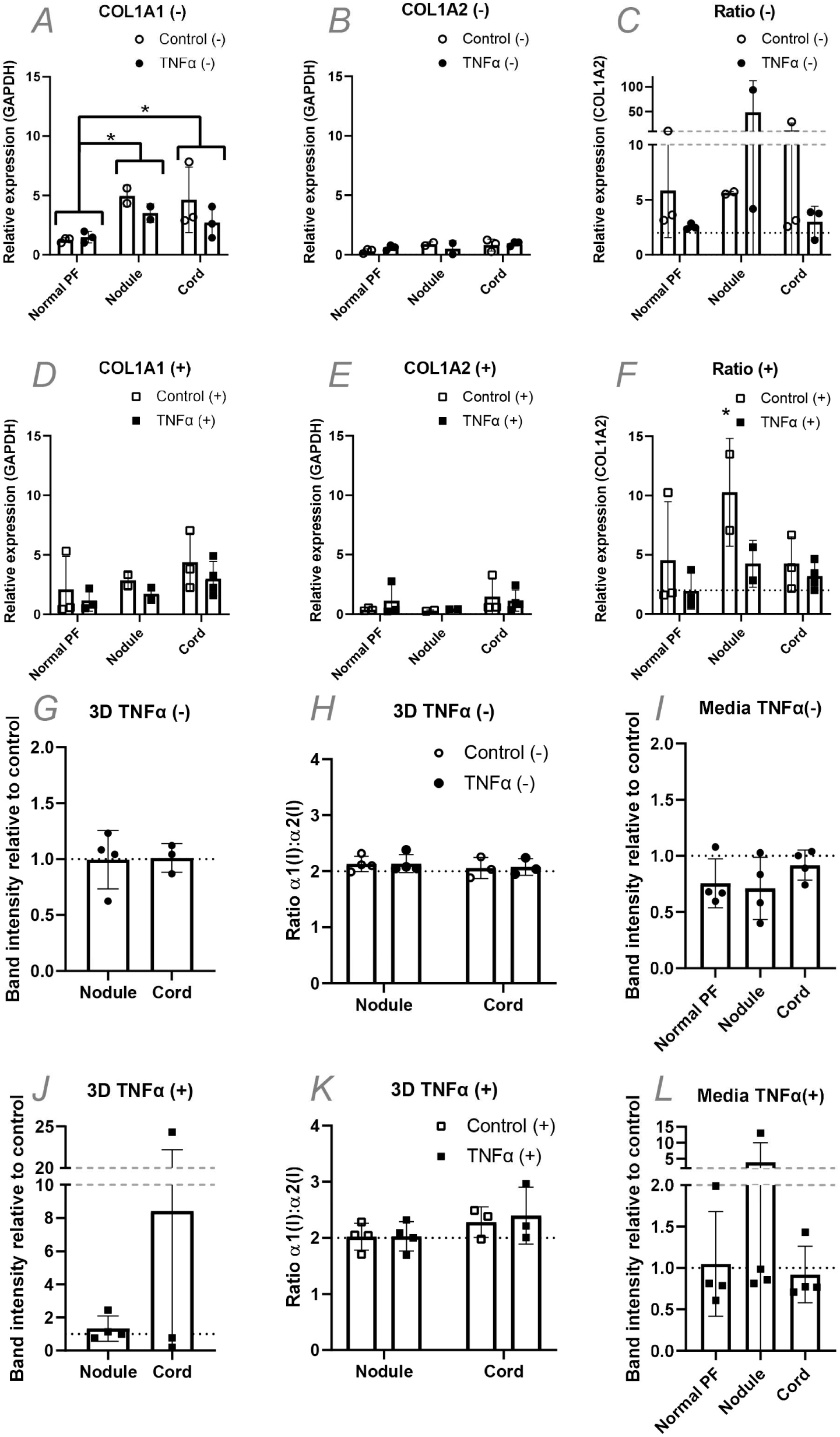
TNF-α has minimal effects on type I collagen synthesis in 3D tendon-like constructs derived from normal PF, Dupuytren’s nodule and Dupuytren’s cord cells. A-F: Analysis of COL1A1 (A, D), COL1A2 (B, D) and COL1A1:COL1A2 (C, F) gene expression by qRT-PCR in 3D tendon-like constructs (n=4) after TNF-α treatment in serum free conditions (-) (A-C) or in the presence of 10% FCS (+) (D-F). G-L: Analysis of type I collagen protein synthesis (G, J), type I collagen homotrimer formation (H, K) and collagen secretion (I, L) by [^14^C]proline labelling in 3D tendon-like constructs after TNF-α treatment in serum free conditions (-) (G-I) or in the presence of 10% FCS (+) (J-L). * p<0.05. Sample details are given in Supplementary Table 3.

TNF-α had no discernible effect on type I collagen protein synthesis, homotrimer production or on newly synthesised collagens in cell culture media in the absence or presence of serum (Fig. 6 G-I). There were no significant differences between cell types or with TNF-α treatment relative to controls for the total α1(I)+α2(I) band intensity (Fig. 6 G&J), representing fully processed type I collagen within 3D tendon-like constructs, or for labelled type I collagen procollagen and processing intermediates secreted into conditioned media (Fig. 6 I&L). Unlike for 2D cultures, TNF-α did not have opposing effects on labelled collagen present in media in the absence or presence of serum (Fig. S2) The α1(I):α2(I) ratio for fully processed type I collagen extracted from 3D tendon-like constructs was not significantly different between conditions, or cell types and was not significantly greater than 2 (Fig. 6 I&L), indicating the 3D tendon-like constructs primarily synthesised heterotrimeric rather than homotrimeric type I collagen.

### Anti-TNF did not reduce type I collagen synthesis in 3D tendon-like constructs or type I collagen homotrimer synthesis in Dupuytren’s tissue

To determine if anti-TNF-α would reduce type I collagen synthesis or processing in 3D tendon-like constructs, the antibody was added to fully formed constructs in the absence of serum. Anti-TNF-α had no effect on the total α1(I)+α2(I) band intensity, representing fully processed type I collagen within and synthesised by 3D tendon-like constructs, as compared to IgG controls (Fig. 7A). The newly synthesised collagens in cell culture media were similarly unaffected by anti-TNF-α treatment, as compared to IgG controls and there were no significant differences between cell types (Fig. 7B). As observed for TNF-α treatment and controls (Fig. 6 H&K) the α1(I):α2(I) ratio for fully processed type I collagen extracted from 3D tendon-like constructs was not significantly greater than 2, changed with anti-TNF-α treatment, or different between cell types.

**Figure 7.**
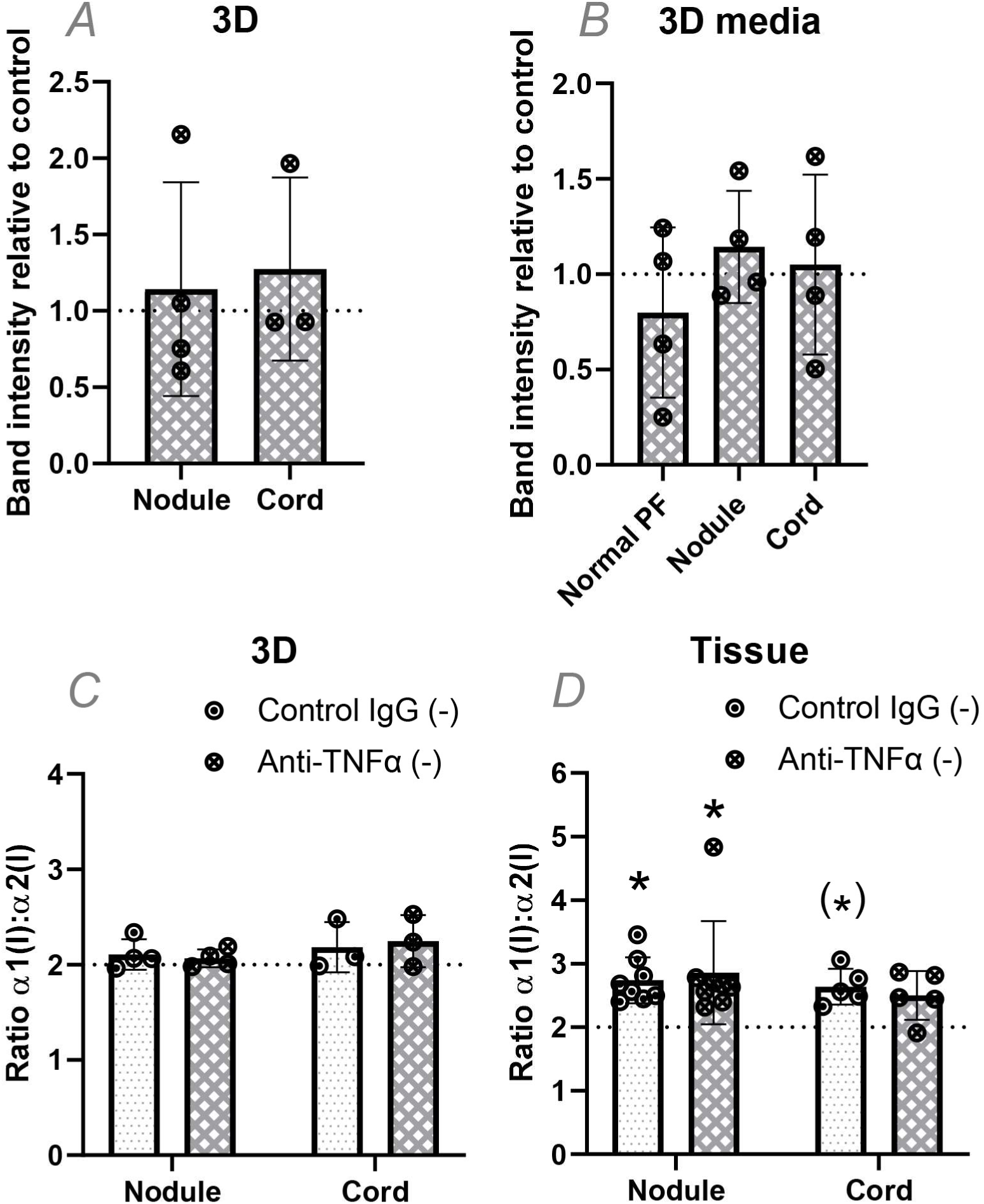
Anti TNF-α does not prevent type I collagen homotrimer formation in Dupuytren’s tissue explants. A-C: Analysis of type I collagen protein synthesis (A), collagen secretion (B) and type I collagen homotrimer formation (C) by [^14^C]proline labelling in 3D tendon-like constructs after control IgG or anti-TNF-α treatment in serum free conditions (-). D: Analysis of type I collagen homotrimer formation by [^14^C]proline labelling in Dupuytren’s nodule or cord tissue explants after treatment with control IgG or with anti-TNF-α in serum free conditions (-). * p<0.05 (1 sample t-tests against a reference value of 2), (*) significant only in a one-tailed test. Sample details are given in Supplementary Table 3.

Unlike for 3D tendon-like constructs, Dupuytren’s tissue explants were found to synthesise homotrimeric type I collagen (Fig. 1 E&F). Nodule and cord tissue explants from Dupuytren’s patients (Supplementary Table 3) were therefore treated with control IgG or anti-TNF-α to determine if anti-TNF would reduce active synthesis of type I collagen homotrimer. For these experiments the total or secreted collagens were not compared due to variability in the size of tissue explants between conditions. The α1(I):α2(I) ratio was significantly greater than 2 for nodule explants (p=0.010 and 0.008 for control and treatment respectively) indicative of type I collagen homotrimer synthesis (Fig. 7D). The cord control appeared significantly greater than 2 only when using a 1-tailed test (p= 0.043). There were no significant differences between anti-TNF-α and control IgG treatment indicating that anti-TNF-α did not reduce active synthesis of type I collagen homotrimer.

## Discussion

In this study, we showed increased COL1A1 and COL1A2 mRNA as well as an increased COL1A1:COL1A2 mRNA ratio in Dupuytren’s tissue as compared to normal PF (Fig. 1 A-C). The results presented are in accordance with previous studies which showed increased COL1A1 mRNA in Dupuytren’s tissue compared to unaffected transverse ligament of the palmar aponeurosis (Ten Dam et al., 2016) and increased COL1A1 but not COL1A2 mRNA in Dupuytren’s tissue, as compared to shoulder capsule (Kilian et al., 2007). For surgical waste human tissue samples we did not distinguish between nodule and cord for RNA analysis, although a prior study found both COL1A1 and COL1A2 mRNA was higher in nodule as compared to cord, despite lower type I collagen protein being present in nodule (van Beuge et al., 2016).

One previous study indicated the presence of type I collagen homotrimer in Dupuytren’s tissue with an increase from 5% in normal PF, to 9% in Dupuytren’s utilising pooled samples (Ehrlich et al., 1982). Utilising metabolic labelling with ^14^C proline, we found a consistent increase in the α1(I):α2(I) polypeptide chain ratio for newly synthesised collagen in Dupuytren’s tissue samples, but noted a skewed distribution with some samples showing a particularly high α1(I):α2(I) chain ratio (Fig. 1 E-F). We hypothesised that individual variation or samples analysed at different stages of disease progression may be responsible for the relative proportion of newly synthesised collagen however, no association with demographics data or disease stage was observed (Fig. 2). It is possible that the current study is underpowered to accurately identify factors or interactions that may direct high type I collagen homotrimer synthesis or that the granularity of the demographics data is insufficient. The mean percentage of homotrimeric collagen is higher than that reported by Ehrlich et. al, although there is a cluster of several samples apparently synthesizing type I collagen homotrimer to a similar extent, at around 10%. Notably our study also provides a snapshot of type I collagen homotrimer synthesis whereas Ehrlich et. al. analysed accumulated homotrimeric type I collagen.

We noted an absence of detectable type I collagen protein synthesis in normal PF by radiolabelling (Fig. 1D) obtaining similar results for healthy canine cranial cruciate ligament and post-natal equine superficial digital flexor tendon. These findings indicate that the type I collagen synthesis detected in Dupuytren’s and ruptured cruciate ligament explants is not due to an acute injury response to dissection. In ruptured canine cranial cruciate ligament (CCL), expression of COL1A1 and COL1A2 mRNA was increased (Fig. 3 A-B). A previous study in canine CCL indicated a significant increase in COL1A2 mRNA, but not of COL1A1 in ruptured ligaments (Clements et al., 2008), whilst COL1A1 mRNA was found to be increased in ruptured human anterior cruciate ligament by in situ hybridization (Fukui et al., 2001). In the present study, the COL1A1:COL1A2 ratio was significantly higher in ruptured canine CCLs but the ratio did not exceed 2 and type I collagen homotrimer synthesis was not apparent (Fig. 3E). Gene expression of *COL1A1* and *COL1A2* in equine SDFT reduced dramatically after 2 years of age (Fig. 3G) and type I collagen protein synthesis was only detected in fetal tendon (Fig. 3F), consistent with the known high-level type I collagen expression during embryonic tendon development (Robbins and Vogel, 1994). No age-related changes in the COL1A1:COL1A2 mRNA ratio were observed in equine SDFT.

Despite a lack of type I collagen protein synthesis, we noted labelled bands in normal PF (Fig. 3A), canine CCL (Fig. 3E), and SDFT (Fig. 3F) tissue extracts. Attempts to identify the labelled proteins using ^13^C proline labelling, band excision and mass spectrometry, or by Western blotting to candidate collagens were unfortunately not informative. However, given the incorporation of ^14^C proline, we expect that these proteins are collagenous in nature or proline-rich.

Whilst we did not detect a difference in *COL1A1* gene expression in 2D cell cultures with TNF-α treatment in the presence or absence of serum, *COL1A2* gene expression was reduced by TNF-α in the presence of serum (Fig. 4) in accordance with a prior study (Verrecchia et al., 2002). TNF-α had opposing effects on new collagen protein secretion in the presence or absence of serum with a more repressive effect in 10% serum. This may relate to the presence of TGF-β in serum, antagonistic TNF-α and TGF-β signalling and the inhibitory effect of serum a2-macroglobulin on cytokine signalling (Verrecchia et al., 2002). Prior work indicated that TNF-α increased COL1A1 mRNA in palmar fibroblasts, from Dupuytren’s patients though not in Dupuytren’s nonpalmar dermal fibroblasts or in palmar fibroblasts from those unaffected by Dupuytren’s, though it is unclear whether treatments were carried out in the presence of serum.

Adult human tenocytes have been previously found to form 3D tendon-like constructs when seeded into a fibrin gel scaffold (Bayer et al., 2010) although these were younger (29 +/- 7.5 years) than the normal PF and Dupuytren’s cells used in our study (range 57-76 years). Hence cells derived from normal PF retain the capacity to synthesise a new collagenous ECM even up to 69 years of age (Supplementary Table 3). Prior radiolabelling of newly synthesized collagen in 3D tendon-like constructs detected pro-forms as well as fully processed collagen (Kalson et al., 2010) following 1 hour of labelling. In the present study fully-processed collagen was primarily detected within constructs which can be attributed to a 3 hour chase, following overnight labelling, allowing the majority of the newly synthesized labelled collagen to traverse the secretory pathway and be fully proteolytically processed to tropocollagen. In contrast the media contained primarily pro-forms; likely due to dilution of the procollagen substrate and processing enzymes in the liquid environment.

In 3D tendon-like constructs the COL1A1 to COL1A2 ratio was decreased with TNF-α treatment in the presence of serum (Fig. 6) indicating potential differential effects on *COL1A1* and *COL1A2* gene expression in 3D culture. However, this was not reflected in an altered type I collagen a1: a2 protein chain ratio. Unlike in 2D culture no differential effects of TNF-α on collagen protein production were observed in 3D culture, either for secreted protein in the media, or that within the constructs. These findings may reflect residual serum proteins being retained within the fully-formed 3D tendon-like constructs, differences in signalling responses in 3D culture or the presence of a collagenous ECM barrier that could prevent TNF-α from interacting with cells.

In the present study anti-TNF was not able to reduce type I collagen synthesis in 3D tendon-like constructs or their media, or to modulate type I collagen homotrimer synthesis in Dupuytren’s tissue (Fig. 7). This appears in contrast to the reported effect of anti-TNF on *COL1A1* gene expression in 2D Dupuytren’s myofibroblasts (Verjee et al., 2013) and reduced type I procollagen in patient nodules 2 weeks after anti-TNF (adalimumab) injection (Nanchahal et al., 2018). Although we did not test anti-TNF in 2D culture, tests in 3D culture or tissue explants are more relevant to the *in vivo* condition and Nanchahal et al. similarly did not detect reduced COL1A1 expression in injected nodules. Interestingly, *in vivo* type I procollagen (measured with an ELISA for the N-propeptide, as a procollagen proxy) was only reduced with the highest anti-TNF dose (p=0.019), which became available during the dose-ranging study (Nanchahal et al., 2018), whereas total type I collagen was not reduced with anti-TNF treatment which is not unexpected given the relative timescales of treatment and prior disease progression.

Herein we utilised IgG as a control for the anti-TNF treatments, whereas saline injection was used as control for the phase 2a trial (Nanchahal et al., 2018), and the control for prior 2D culture studies is ambiguous (Verjee et al., 2013). The anti-TNF biologic infliximab does not bind soluble or membrane-bound mouse TNF-α yet improves mobility and suppresses joint inflammation, weight loss and pain in mice overexpressing mouse TNF-α (TNF^DARE^) (Naylor et al., 2018). Indeed, Fc receptor-mediated effects contribute to the therapeutic response to infliximab and adalimumab in mouse models of inflammatory bowel disease (IBD) (McRae et al., 2016) and IBD patients only respond to full IgG1 anti-TNF therapies via macrophage IL-10 signalling (Koelink et al., 2020; Levin et al., 2016; Vos et al., 2011). Indeed immune cell populations including macrophages are present in Dupuytren’s disease tissue (Andrew et al., 1991; Verjee et al., 2013). Hence it is possible that the beneficial effects of the anti-TNF adulimumab in Dupuytren’s disease could be Fc mediated and not related to modulation of collagen synthesis.

## Methods

### Sample collection

After full informed written consent (REC13/NW/0352 and REC 14/NW/0162), excised Dupuytren’s tissue (fibrotic palmar fascia) and control non-diseased normal palmar fascia (PF), were taken from human patients undergoing surgery for Dupuytren’s contracture and Carpal tunnel syndrome respectively. Surgery was carried out within Royal Liverpool and Broadgreen University Hospitals NHS Trust (now Liverpool University Hospitals NHS Foundation Trust) or the Warrington and Halton Hospitals NHS Foundation Trust (now Warrington and Halton Teaching Hospitals NHS Foundation Trust). Self-reported demographics were collected by questionnaire, and for the Dupuytren’s patients, disease stage information was collected from clinical records. Equine superficial digital flexor tendon (SDFT) and healthy canine cranial cruciate ligament (CCL) were collected as previously described (Ali et al., 2018; Lee et al., 2018), with local ethical approval (VREC186, VREC62 and RETH00000553). Ruptured CCL, otherwise discarded as clinical waste, was obtained from dogs undergoing stifle/knee stabilisation at the University of Liverpool’s Small Animal Teaching Hospital (VREC63). Surgical samples were reserved in cold sterile saline and processed on the day of collection according to the selected analysis method. During processing of Dupuytren’s samples perinodular fat was first removed and samples separated into cord and nodule where indicated.

### Tissue pulse-chase with ^14^C-L-proline

Tissue samples were aseptically dissected into pieces, with an estimated wet weight of 25-50 mg or volume of ∼25-50 μL each, under PBS or Dulbecco’s minimal essential medium (DMEM), containing penicillin/streptomycin (1% v/v). After a 30-60 minute pre-equilibration at 37°C in DMEM containing penicillin/streptomycin (1% v/v), L-glutamine (2 mM), L-ascorbic acid 2-phosphate (200 μM) and β-aminopropionitrile (400 μM) (labelling media) with ∼1-5 pieces per 1 ml of media, pulse–chase experiments were performed in labelling media supplemented with 2.5 μCi/ml [^14^C]proline (Perkin Elmer, Waltham, USA). After incubation in supplemented labelling media for 18 hrs, tissue samples were transferred to labelling media without [^14^C]proline for 3 hrs. Subsequently, 1-2 tissue pieces were extracted in 100 μl aliquots of salt extraction buffer (1 M NaCl, 25 mM EDTA, 50 mM Tris-HCl, pH 7.4) containing protease inhibitors. Extracts were analysed by electrophoresis on 6% Tris-Glycine gels (ThermoFisher, Waltham, USA) with delayed reduction (Sykes et al., 1976) or on 4% Tris-Glycine gels (Invitrogen/ThermoFisher, discontinued) under reducing conditions. The gels were fixed in 10% methanol, 10% acetic acid, dried under vacuum, and exposed to a phosphorimaging plate (BAS-IP MS). Phosphorimaging plates were processed using a phosphorimager (Typhoon FLA7000 IP, GE Healthcare/Cytiva, Marlborough, USA) and densitometry carried out using ImageQuant (GE Healthcare/Cytiva). The α1(I):α2(I) chain ratio was corrected for the relative amount of proline residues in each chain (240 in α1(I) and 204 in α2(I) such that the ratio was divided by 1.1765) before analysis, and converted to a percentage homotrimeric collagen using the formula; (ratio-2)x100/ratio.

### Cell isolation and culture

Tissue samples were dissected into small pieces and digested with 1-2 mg/ml (275 U/ml) collagenase type II (Worthington, Columbus, USA) in 30 ml Dulbecco’s minimal essential medium (DMEM) containing 1 % (v/v) penicillin/streptomycin and 2% FBS (F7524, Batch 024M3398, Sigma-Aldrich, Missouri, USA), at 37°C overnight in a rotary shaker. Dupuytren’s cell suspensions were strained in a 70 um cell strainer, centrifuged for 10 minutes at 2300 rpm and the cell pellets re-suspended in DMEM with L-glutamine supplemented with 10% FBS, 1 % (v/v) penicillin/streptomycin and 0.5 µg/mL fungizone (Thermo Fisher). Cells were seeded at 0.5-3.0 x 10^5^ cells per cm^2^ in a T25 or T75 flask, grown at 37°C in 5% CO_2_ and cryopreserved at passage 0 or 1. Normal palmar fascia samples were digested with 2 mg/ml collagenase in 15m of media with cell straining omitted for 3 of the 4 samples. Cells were seeded directly into 1 well of a 6-well plate and cryopreserved at passage 1.

Dupuytren’s cells split 1:2 at passage 1 or 2 and normal PF cells at passage 3 or 4, were used to establish 3D tendon-like constructs as described (Kapacee et al., 2008), with the following modifications. Constructs for ultrastructural analysis and radiolabelling were established in 24-well rather than 6-well plates as were those for qRT-PCR, excepting those derived from normal PF cells and cord-derived cells from sample 161012-62-F. To form constructs in 24-well plates (Fig. S3), pins were positioned 5 mm apart, the volume of thrombin to use was titrated in advance for each batch and constructs were seeded with 2.5 x 10^5^ cells per well. For 6-well plates constructs were seeded with 6 x 10^5^ cells per well. Scoring and media changes took place 3 times a week (every 2-3 days).

### Cell and tissue treatments

Prior to treatment the media was changed to that supplemented with non-essential amino acids for 2D cell cultures, L-ascorbic acid 2-phosphate (200 uM), β-aminopropionitrile (400 uM), without serum as indicated or for explants, and cultures incubated at 37°C in 5% CO_2_ for 16-18 hours. The media was then replaced with that containing 10ng/ml human TNF-α (300-01A, Peprotech, London, UK), vehicle control, mouse anti-human TNF-α antibody (MAB610, R&D Systems, Minneapolis, USA) or mouse IgG1 Isotype Control (MAB002), along with 2.5 μCi/ml [^14^C]proline overnight as required. Labelling media was reserved and chase media without supplements added for 3 hours. Cells were used at passage 3 or 4 and at 80% confluence, constructs were fully contracted and tissue samples were treated on the day of collection. Labelled tissues and 3D constructs were snap frozen and labelled collagen extracted with salt-extraction buffer. Salt extracts and labelling media were analysed by delayed reduction electrophoresis as described above.

### Quantitative real-time PCR (qRT-PCR)

Tissue samples and 3D tendon-like constructs were immersed in RNAlater (Qiagen, Hilden, Germany) and stored at -20°C until analysis. The tissue was homogenised with a Mikro-Dismembrator (Sartorius, Göttingen, Germany) or a pestle and mortar in 1 ml of TRI Reagent® (Sigma-Aldrich, Poole, UK). 0.1 ml 1-bromo-3-chloropropane (BCP) (Sigma-Aldrich) was added to the homogenate and centrifuged for 15 minutes at 12,000 x g at 4°C. 1 μg of total RNA was reverse transcribed with M-MLV reverse transcriptase using random primers according to the manufacturer’s protocol (Promega, Madison, USA). qRT-PCR was performed in a 25 μl reaction volume containing; complementary DNA (cDNA) (5 ng), primers designed for the gene of interest (Table 1) and GoTaq(R) qPCR Master Mix (Promega). Samples were run on an AB 7300 Real Time PCR System (Applied Biosystems, Waltham, USA) using the following amplification conditions; 2 minutes at 95°C followed by 40 cycles of 15 seconds at 95°C and 1 minute at 60°C. Gene expression was calculated relative to GAPDH, which was determined to be a suitable reference gene after assessing its stability using the geNorm method (Vandesompele et al., 2002).

**Table 1:**
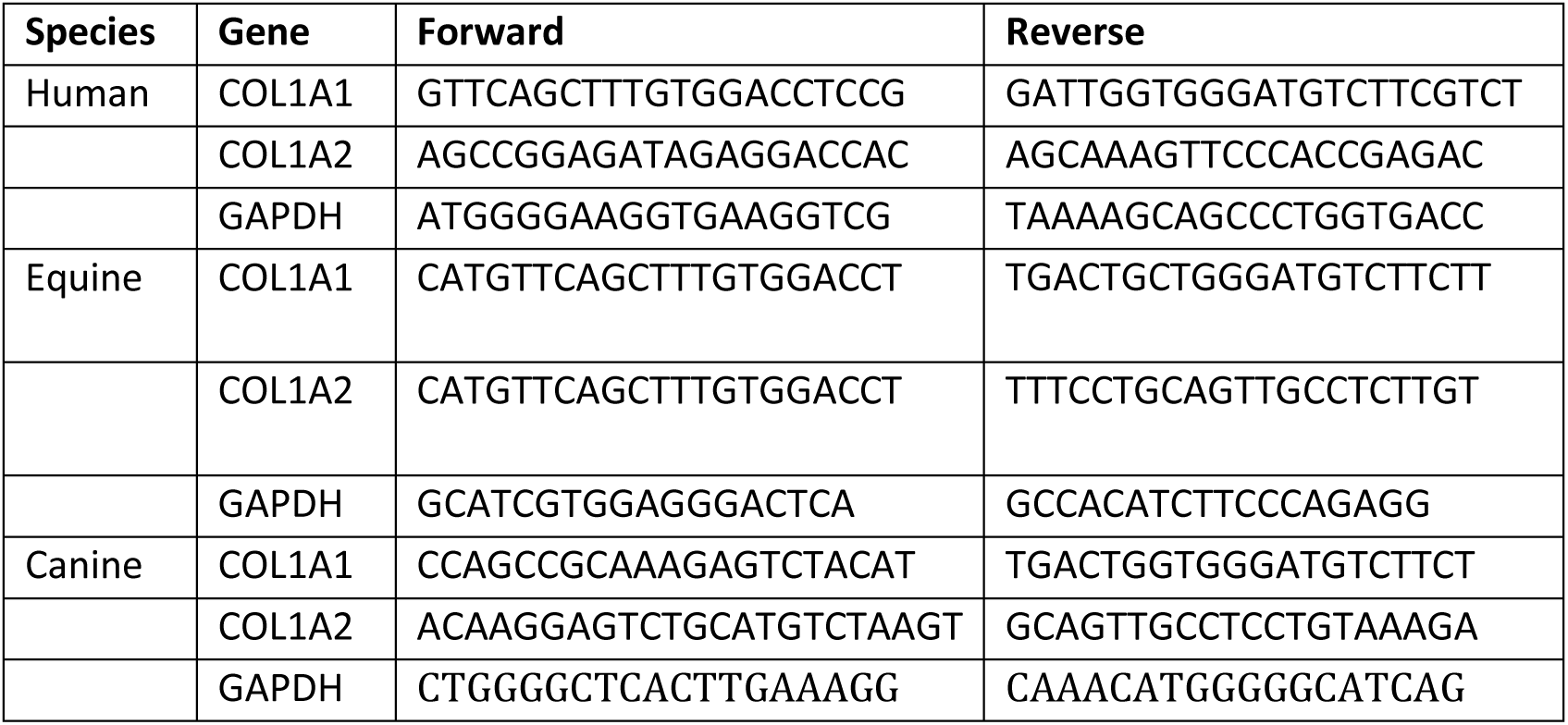
Primer Sequences used for qPR-PCR.

RNA was first extracted from cultured cells using Trizol. An equal volume of 100% ethanol was added to the aqueous phase after phase separation with chloroform and then loaded onto an RNeasy column (Qiagen). RNA was purified following the manufacturer’s protocol with the following modifications: 30 s rather than 15 s spins, only one wash with buffer RPE, an optional spin to exclude RPE buffer carryover was included and the final elution was carried out 1-2 min after adding RNase-free water. Constructs were homogenised using a steel ball lysing matrix and a FastPrep 24 tissue homogeniser (MP Biomedicals, Santa Ana, CA, USA). RNA was extracted from homogenised samples using a RNeasy kit (Qiagen) as per the manufacturer’s instructions. cDNA was synthesised in a 25 µl reaction from 0.5-1 ug total RNA for cells and 0.15 ug for constructs as described. cDNA synthesis and qPCR were performed as described (Lee et al., 2022) using human primers shown in Table 1.

### Transmission electron microscopy (TEM)

TEM was carried out as described (Kharaz et al., 2016) but modified for constructs derived from normal PF cells, for which an initial fixation (∼5 mins) was carried out in 4% paraformaldehyde in PBS before transferring half the fixed construct to 2.5% glutaraldehyde in 0.1M cacodylate. Semi-thin (0.5 um) sections were stained with 1% toluidine blue and visualised by light microscopy.

### Statistical analysis

Data analysis was performed using SigmaPlot Version 14.0 (Systat, San Jose, USA) and graphing carried out with GraphPad Prism 8 for Windows (GraphPad Software, La Jolla California USA, www.graphpad.com), unless otherwise indicated. Plots show individual data points together with the mean and +/- 1 SD unless otherwise indicated. Following a log_10_ transformation as required, COL1 gene expression data and sample ages were analysed using a Welch’s t-test. Polypeptide chain ratios were transformed with an inverse function and analysed using a one-sample t-test. qPCR and collagen synthesis data were analysed by 1-way or 2-way ANOVA or by a one-sample t-test as appropriate. A prior suitable Box-Cox transformation was carried out if assumptions of normality and equal variance were not met. PCA and associated graphing was carried out using Minitab® Statistical Software Version 18 (State College, PA: Minitab, Inc. (www.minitab.com)) as was Box-Cox data transformation. For PCA incorporating demographic data, self-reported textual data was coded in a binary manner except for diabetes - where diet-controlled (coded 1) was distinguished from type I (coded 2), smoking - where previous smokers were coded 1 and current smokers coded 2, alcohol consumption - where those consuming below the recommended limit were coded 1 and those above coded 2, and exercise - which was coded from 0-3 based on apparent intensity/duration.

## Author contributions

Conceptualization: EGC-L. Funding acquisition: EGC-L. Resources: GCh, DB, RP, EJC, PDC. Methodology: KAW, GCo, KJL, EGC-L. Investigation: KAW, KJL, EB, AC, JAG. Data curation: KAW, KJL, EB, EGC-L. Project administration: RP, EJC, PDC, EGC-L. Supervision: EJC, PDC, EGC-L. Formal Analysis: KAW, KJL, EB, EGC-L. Visualization: KAW, KJL, EB, EGC-L. Writing – original draft: EGC-L. Writing – review & editing: KAW, KJL, EB, JAG, GC, GCh, DB, RP, EJC, PDC, EGC-L.

## Acknowledgements

The assistance of the surgical teams and research nurses at the Royal Liverpool and Broadgreen University Hospitals NHS Trust and the Warrington and Halton Hospitals NHS Foundation Trust, as well as the contribution of the patients donating samples for the study is gratefully acknowledged. The authors would like to thank the staff at the Philip Leverhulme Equine Hospital and the Small Animal Teaching Hospital, as well as the owners that donated tissue for use in research. The invaluable assistance of Dr Adam Janvier in maintaining 3D constructs, as well as Mr Adrian Chojnowski at the Norfolk and Norwich University Hospitals NHS Foundation Trust, and Dr Graham Riley and Professor Ian Clark at the University of East Anglia in providing human surgical samples (REC 10/H0310/03) early in the study is gratefully acknowledged.

This study was funded by the UK Medical Research Council (MRC) (MR/J002909). KJL was supported by the Marjorie Forrest Bequest and by the Institute of Ageing and Chronic Disease at the University of Liverpool then MRC (MR/R00319X/1). EB was supported by Versus Arthritis (21809). JAG was supported by The Medical Research Council Versus Arthritis Centre for Integrated Research into Musculoskeletal Ageing (CIMA).

## Supplementary Figures

**Figure S1:**
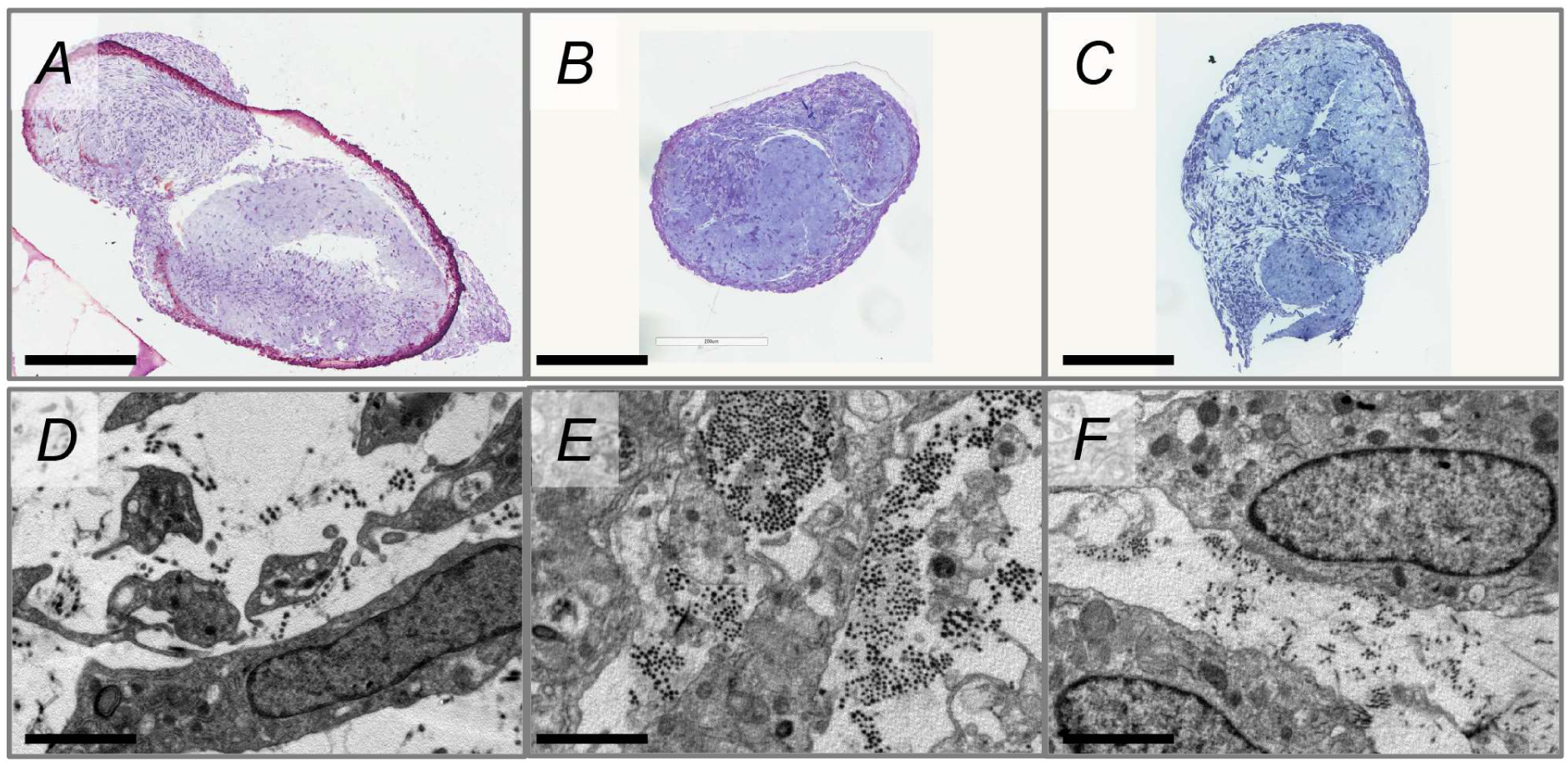
*De novo* collagen fibrils are present within tendon-like structures in 3D culture of normal palmar fascia and Dupuytren’s cells. A-C: Semi-thin toluidine blue-stained sections of 3D tendon-like constructs (bar 200 um). D-F: Transmission electron microscopy images of 3D tendon-like constructs (bar 2 um). Constructs were derived from normal palmar fascia cells (A&D), Dupuytren’s nodule (B&E) or Dupuytren’s cord (C&F). Sample IDs are shown in Supplementary Table 3.

**Figure S2.**
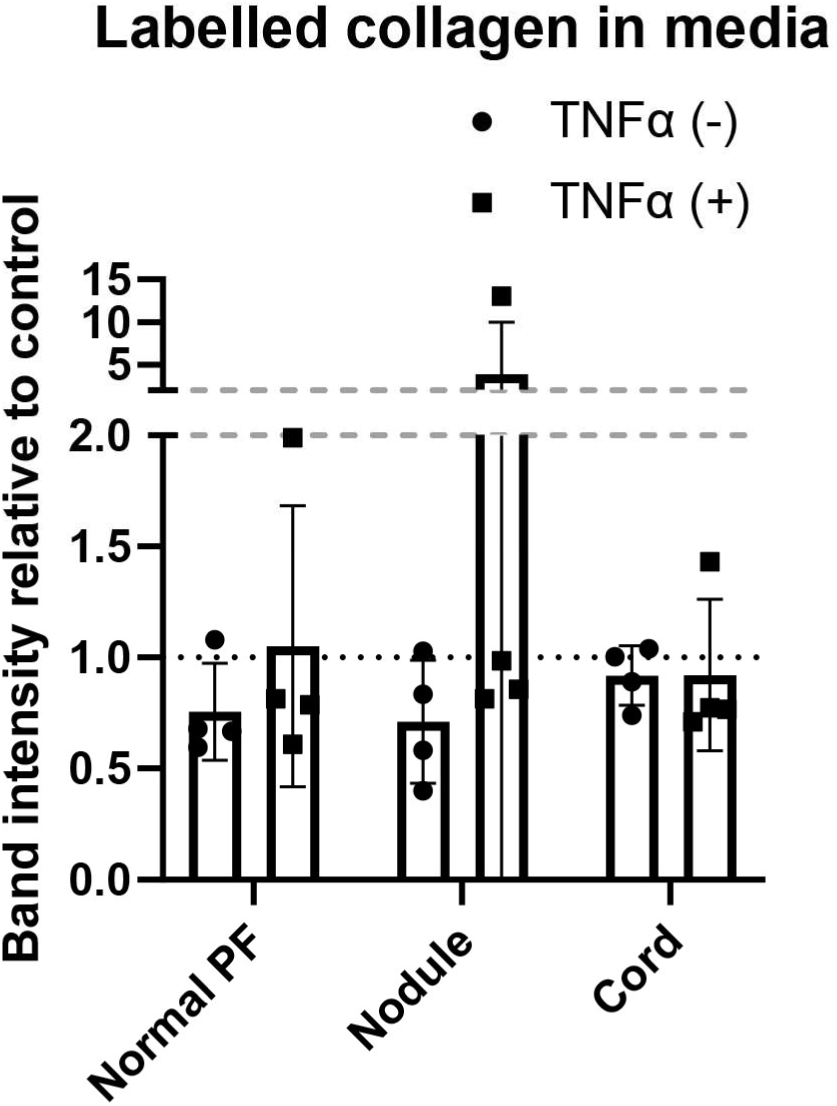
Comparison of the amount of labelled collagen in the media of tendon-like constructs treated with TNF-α in the absence or presence of serum. Densitometric quantification of the relative amounts of radiolabelled (pro)collagen present in conditioned media from normal PF, Dupuytren’s nodule and Dupuytren’s cord cells (n=4) after TNF-α treatment, as compared to control treatments, in serum free conditions (-) and in the presence of 10% FCS (+).

**Figure S3.**
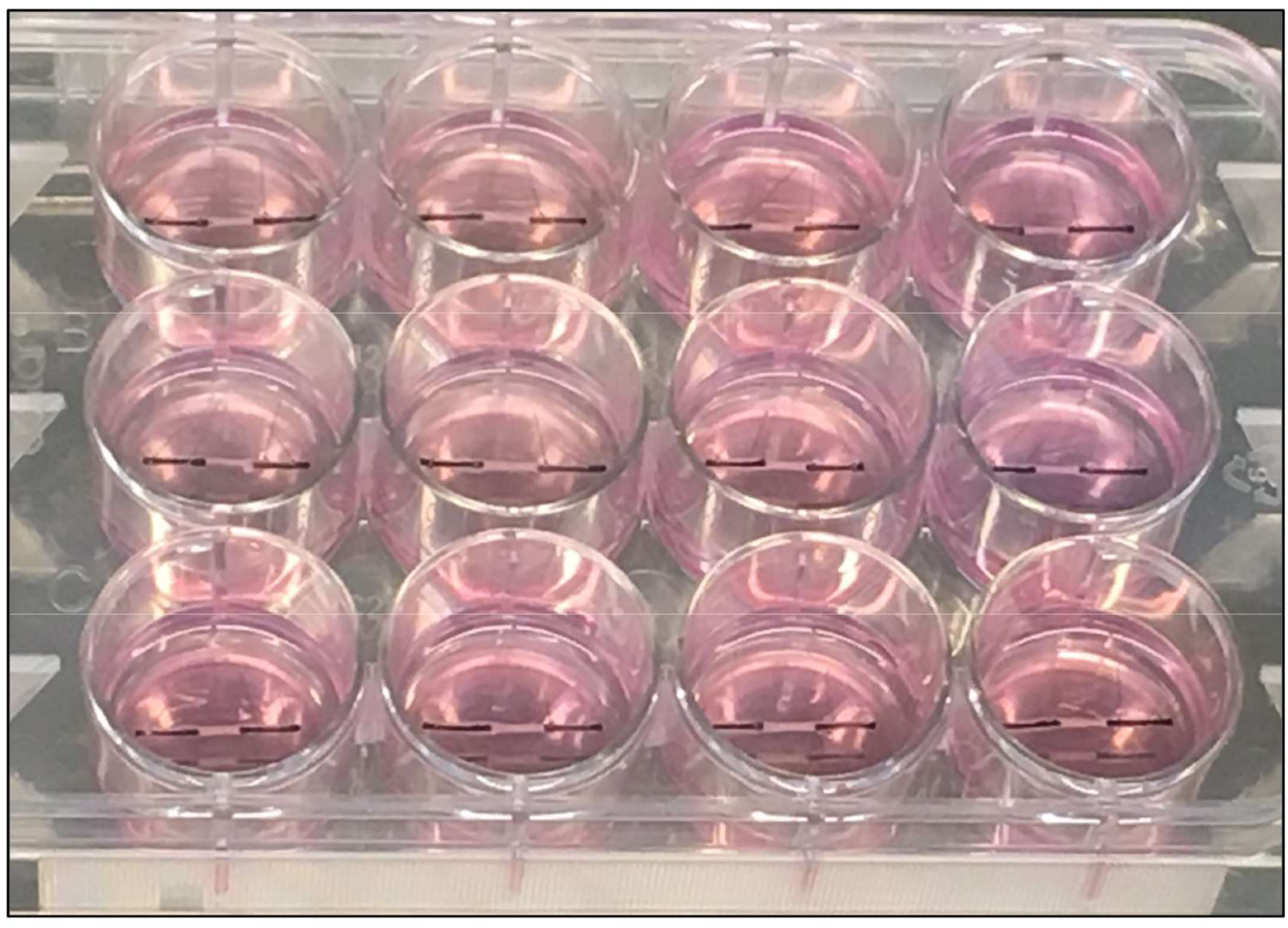
3D tendon-like constructs formed in a 24 well plate. Image of a 24-well plate containing fully-formed 3D tendon-like constructs between pinned sutures (black) in each well.

## Supplementary Tables

**Supplementary Table 1:**
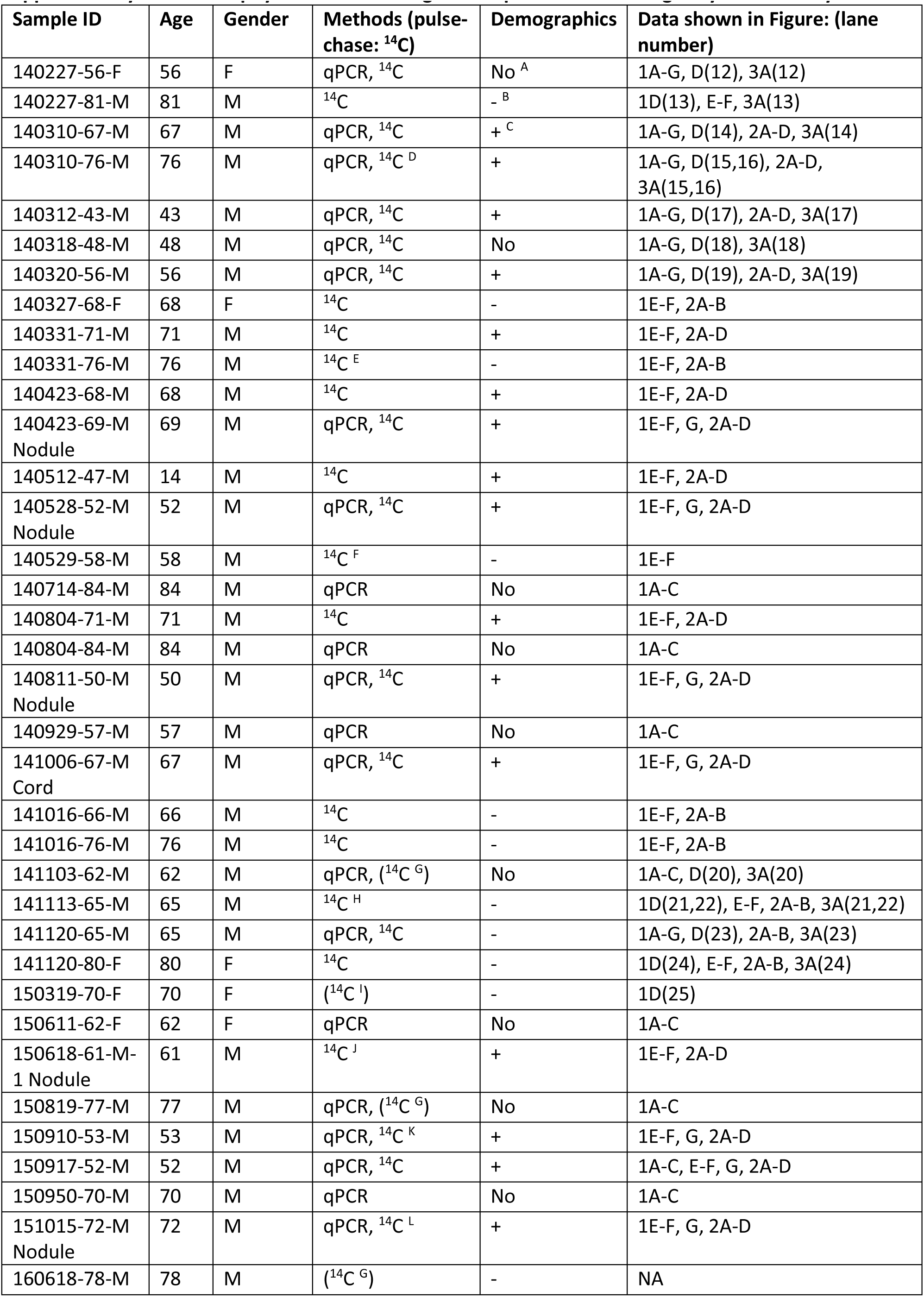
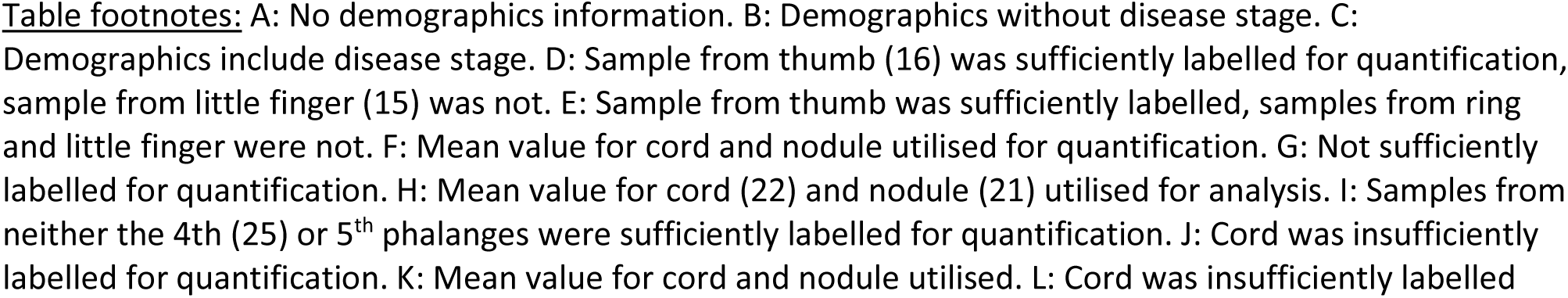
Dupuytren’s human surgical samples used for collagen synthesis assays.

**Supplementary Table 2:** Human normal Palmar Fascia (PF) surgical samples used for collagen synthesis assays.

| Sample ID | Age | Gender | Methods (pulse-chase: $^{14}\text{C}$) | Data shown in Figure: (lane number) |
| --- | --- | --- | --- | --- |
| 140723-82-F | 82 | F | qPCR, $^{14}\text{C}$ | 1A-C, D(4), 3A(4) |
| 140821-38-M | 38 | M | qPCR, $^{14}\text{C}$ | 1A-C, D(2), 3A(2) |
| 140821-39-F | 39 | F | qPCR, $^{14}\text{C}$ | 1A-C, 3A(11) |
| 140822-54-F | 54 | F | qPCR, $^{14}\text{C}$ | 1A-C, D(3), 3A(3) |
| 140822-55-F | 55 | F | qPCR, $^{14}\text{C}$ | 1A-C, D(1), 3A(1) |
| 140822-68-F | 68 | F | qPCR, $^{14}\text{C}$ | 1A-C, D(6), 3A(6) |
| 140911-48-M | 48 | M | $^{14}\text{C}$ | 1D(10), 3A(10) |
| 141002-44-F | 44 | F | qPCR | 1A-C, D(7), 3A(7) |
| 141003-79-F | 79 | F | qPCR | 1A-C |
| 141007-57-F | 57 | F | qPCR | 1A-C |
| 141009-25-F | 25 | F | $^{14}\text{C}$ | 1D(8) |
| 141013-34-M | 34 | M | qPCR | 1A-C |
| 141013-67-M | 67 | M | qPCR | 1A-C |
| 141103-51-F | 51 | F | $^{14}\text{C}$ | 1D(9) |
| 141125-77-F | 77 | F | $^{14}\text{C}$ | 1D(5) |
| 141128-50-F | 50 | F | qPCR | 1A-C |
| 141218-53-M | 53 | M | qPCR | 1A-C |

**Supplementary Table 3.** Human surgical samples used for cell culture and treatments.

| Sample ID | Age | Gender | Tissue type | Data shown in Figure: |
| --- | --- | --- | --- | --- |
| 161012-62-F | 62 | F | Dupuytren's Cord & Nodule | 4, 5 E& H(nodule), F,I(cord), 6 A-F(cord), G-H & J-K(nodule), I, L, 7 A&C(nodule), 7B |
| 161109-70-M | 70 | M | Dupuytren's Cord & Nodule | 4, 5B (nodule, 6-11), 6 A-F(cord), G-L, 7 A-C |
| 161130-65-M | 65 | M | Dupuytren's Cord & Nodule | 4, 5C (nodule, 12-17), 6 A-F(nodule), G-L, 7A-C |
| 161130-76-M | 76 | M | Dupuytren's Cord & Nodule | 4, 5A (cord, 1-4), 6 A-L, S1 B&E(nodule), D&F(cord), 7 A-C |
| 170201-67-M | 67 | M | Normal Palmar Fascia | 4, 6 A-F, I, L, 7B |
| 170206-69-M | 69 | M | Normal Palmar Fascia | 4, 6 I, L, 7B |
| 170301-61-F | 61 | F | Normal Palmar Fascia | 4, 5, 6 A-F, S1A, D, I, L, 7B |
| 170313-57-F | 57 | F | Normal Palmar Fascia | 4, 5 D&G, 6 A-F, I, L, 7B |
| 170403-68-M | 68 | M | Dupuytren's Cord & Nodule | 7D |
| 170403-86-M | 86 | M | Dupuytren's Cord & Nodule | 7D |
| 170412-51-M | 51 | M | Dupuytren's Nodule | 7D |
| 170412-68-F | 68 | F | Dupuytren's Nodule | 7D |
| 170508-76-M | 76 | M | Dupuytren's Cord & Nodule | 7D |
| 170515-65-M | 65 | M | Dupuytren's Cord & Nodule | 7D |
| 170802-80-M | 80 | M | Dupuytren's Nodule | 7D |
| 170816-73-F | 73 | F | Dupuytren's Cord | 7D |
| 170816-78-M | 78 | M | Dupuytren's Nodule | 7D |

## Notes

**Conflict-of-interest statement:** KJL has received income as salary from Convatec Inc and GC from BOLDSCIENCE Inc. DB acts as a paid consultant for both Acumed LLC (Hilsboro, OR, USA) and Swemac Innovations (Linkoping, Sweden) and is paid in accordance with industry guidelines for consultancy, teaching and product development. DB received no payment for anything associated with this manuscript.

The other authors have declared that no conflict of interest exists.

### Competing Interest Statement

KJL has received income as salary from Convatec Inc and GC from BOLDSCIENCE Inc. DB acts as a paid consultant for both Acumed LLC (Hilsboro, OR, USA) and Swemac Innovations (Linkoping, Sweden) and is paid in accordance with industry guidelines for consultancy, teaching and product development. DB received no payment for anything associated with this manuscript. The other authors have declared that no conflict of interest exists.

### Summary of Updates

Data on TNF-alpha added and proteomics data removed (to be incorporated into a sister manuscript with additional data). Manuscript text and figures adjusted accordingly.

## References

Ali, O.J., E.J. Comerford, P.D. Clegg, and E.G. Canty-Laird. 2018. Variations during ageing in the three-dimensional anatomical arrangement of fascicles within the equine superficial digital flexor tendon. Eur Cell Mater. 35:87–102.

Andrew, J.G., S.M. Andrew, A. Ash, and B. Turner. 1991. An investigation into the role of inflammatory cells in Dupuytren’s disease. J Hand Surg Br. 16:267–271.

Bayer, M.L., P. Schjerling, A. Herchenhan, C. Zeltz, K.M. Heinemeier, L. Christensen, M. Krogsgaard, D. Gullberg, and M. Kjaer. 2014. Release of tensile strain on engineered human tendon tissue disturbs cell adhesions, changes matrix architecture, and induces an inflammatory phenotype. PLoS One. 9:e86078.

Bayer, M.L., C.Y. Yeung, K.E. Kadler, K. Qvortrup, K. Baar, R.B. Svensson, S.P. Magnusson, M. Krogsgaard, M. Koch, and M. Kjaer. 2010. The initiation of embryonic-like collagen fibrillogenesis by adult human tendon fibroblasts when cultured under tension. Biomaterials. 31:4889–4897.

Brickley-Parsons, D., M.J. Glimcher, R.J. Smith, R. Albin, and J.P. Adams. 1981. Biochemical changes in the collagen of the palmar fascia in patients with Dupuytren’s disease. J Bone Joint Surg Am. 63:787–797.

Canty, E.G., and K.E. Kadler. 2005. Procollagen trafficking, processing and fibrillogenesis. J Cell Sci. 118:1341–1353.

Clements, D.N., S.D. Carter, J.F. Innes, W.E. Ollier, and P.J. Day. 2008. Gene expression profiling of normal and ruptured canine anterior cruciate ligaments. Osteoarthritis Cartilage. 16:195–203.

Craxford, S., and P.G. Russell. 2016. Dupuytren’s disease. Surgery (Oxford). 34:139–143.

Dibenedetti, D.B., D. Nguyen, L. Zografos, R. Ziemiecki, and X. Zhou. 2011. Prevalence, incidence, and treatments of Dupuytren’s disease in the United States: results from a population-based study. Hand. 6:149–158.

Ehrlich, H.P., H. Brown, and B.S. White. 1982. Evidence for type V and I trimer collagens in Dupuytren’s Contracture palmar fascia. Biochem Med. 28:273–284.

Forrester, H.B., P. Temple-Smith, S. Ham, D. de Kretser, G. Southwick, and C.N. Sprung. 2013. Genome-wide analysis using exon arrays demonstrates an important role for expression of extra-cellular matrix, fibrotic control and tissue remodelling genes in Dupuytren’s disease. PLoS One. 8:e59056.

Fukui, N., Y. Katsuragawa, A. Kawakami, H. Sakai, H. Oda, and K. Nakamura. 2001. Metabolic activity in disrupted human anterior cruciate ligament. Evaluation of procollagen mRNA expression in 29 patients. Joint Bone Spine. 68:318-326.

Grazina, R., S. Teixeira, R. Ramos, H. Sousa, A. Ferreira, and R. Lemos. 2019. Dupuytren’s disease: where do we stand? EFORT Open Rev. 4:63–69.

Hindocha, S., D.A. McGrouther, and A. Bayat. 2009. Epidemiological evaluation of Dupuytren’s disease incidence and prevalence rates in relation to etiology. Hand. 4:256–269.

Hindocha, S., J.K. Stanley, J.S. Watson, and A. Bayat. 2008. Revised Tubiana’s staging system for assessment of disease severity in Dupuytren’s disease-preliminary clinical findings. Hand. 3:80–86.

Janvier, A.J., E. Canty-Laird, and J.R. Henstock. 2020. A universal multi-platform 3D printed bioreactor chamber for tendon tissue engineering. J Tissue Eng. 11:2041731420942462.

Jung, J., G.W. Kim, B. Lee, J.W.J. Joo, and W. Jang. 2019. Integrative genomic and transcriptomic analysis of genetic markers in Dupuytren’s disease. BMC Med Genomics. 12:98.

Kalson, N.S., D.F. Holmes, Z. Kapacee, I. Otermin, Y. Lu, R.A. Ennos, E.G. Canty-Laird, and K.E. Kadler. 2010. An experimental model for studying the biomechanics of embryonic tendon: Evidence that the development of mechanical properties depends on the actinomyosin machinery. Matrix Biol. 29:678–689.

Kapacee, Z., S.H. Richardson, Y. Lu, T. Starborg, D.F. Holmes, K. Baar, and K.E. Kadler. 2008. Tension is required for fibripositor formation. Matrix Biol. 27:371–375.

Kapacee, Z., C.-Y.C. Yeung, Y. Lu, D. Crabtree, D.F. Holmes, and K.E. Kadler. 2010. Synthesis of embryonic tendon-like tissue by human marrow stromal/mesenchymal stem cells requires a three-dimensional environment and transforming growth factor [beta]3. Matrix Biology. 29:668–677.

Kharaz, Y.A., S.R. Tew, M. Peffers, E.G. Canty-Laird, and E. Comerford. 2016. Proteomic differences between native and tissue-engineered tendon and ligament. Proteomics. 16:1547–1556.

Kilian, O., U. Pfeil, S. Wenisch, C. Heiss, R. Kraus, and R. Schnettler. 2007. Enhanced alpha 1(I) mRNA expression in frozen shoulder and dupuytren tissue. Eur J Med Res. 12:585–590.

Koelink, P.J., F.M. Bloemendaal, B. Li, L. Westera, E.W.M. Vogels, M. van Roest, A.K. Gloudemans, A.B. van ’t Wout, H. Korf, S. Vermeire, A.A. Te Velde, C.Y. Ponsioen, G.R. D’Haens, J.S. Verbeek, T.L. Geiger, M.E. Wildenberg, and G.R. van den Brink. 2020. Anti-TNF therapy in IBD exerts its therapeutic effect through macrophage IL-10 signalling. Gut. 69:1053-1063.

Krefter, C., M. Marks, S. Hensler, D.B. Herren, and M. Calcagni. 2017. Complications after treating Dupuytren’s disease. A systematic literature review. Hand Surg Rehabil. 36:322–329.

Lam, W.L., J.M. Rawlins, R.O. Karoo, I. Naylor, and D.T. Sharpe. 2010. Re-visiting Luck’s classification: a histological analysis of Dupuytren’s disease. J Hand Surg Eur Vol. 35:312–317.

Layton, T., and J. Nanchahal. 2019. Recent advances in the understanding of Dupuytren’s disease. F1000Res. 8.

Lee, K.J., P.D. Clegg, E.J. Comerford, and E.G. Canty-Laird. 2018. A comparison of the stem cell characteristics of murine tenocytes and tendon-derived stem cells. BMC Musculoskelet Disord. 19:116.

Lee, K.J., L. Rambault, G. Bou-Gharios, P.D. Clegg, R. Akhtar, G. Czanner, R. van ‘t Hof, and E.G. Canty-Laird. 2022. Collagen (I) homotrimer potentiates the osteogenesis imperfecta (oim) mutant allele and reduces survival in male mice. Disease Models & Mechanisms. 15:dmm049428.

Levin, A.D., M.E. Wildenberg, and G.R. van den Brink. 2016. Mechanism of Action of Anti-TNF Therapy in Inflammatory Bowel Disease. J Crohns Colitis. 10:989–997.

Luck, J.V. 1959. Dupuytren’s contracture; a new concept of the pathogenesis correlated with surgical management. J Bone Joint Surg Am. 41-A:635-664.

Makareeva, E., S. Han, J.C. Vera, D.L. Sackett, K. Holmbeck, C.L. Phillips, R. Visse, H. Nagase, and S. Leikin. 2010. Carcinomas Contain a Matrix Metalloproteinase–Resistant Isoform of Type I Collagen Exerting Selective Support to Invasion. Cancer Res. 70:4366–4374.

McRae, B.L., A.D. Levin, M.E. Wildenberg, P.J. Koelink, P. Bousquet, I. Mikaelian, A.S. Sterman, S. Bryant, G. D’Haens, R. Kamath, J. Salfeld, and G.R. van den Brink. 2016. Fc Receptor-mediated Effector Function Contributes to the Therapeutic Response of Anti-TNF Monoclonal Antibodies in a Mouse Model of Inflammatory Bowel Disease. J Crohns Colitis. 10:69–76.

Mori, K., A. Hatamochi, H. Ueki, A. Olsen, and S.A. Jimenez. 1996. The transcription of human α1(I) procollagen gene (COL1A1) is suppressed by tumour necrosis factor-α through proximal short promoter elements: evidence for suppression mechanisms mediated by two nuclear-factorbinding sites. Biochemical Journal. 319:811–816.

Nanchahal, J., C. Ball, D. Davidson, L. Williams, W. Sones, F.E. McCann, M. Cabrita, J. Swettenham, N.J. Cahoon, B. Copsey, E. Anne Francis, P.C. Taylor, J. Black, V.S. Barber, S. Dutton, M. Feldmann, and S.E. Lamb. 2018. Anti-Tumour Necrosis Factor Therapy for Dupuytren’s Disease: A Randomised Dose Response Proof of Concept Phase 2a Clinical Trial. EBioMedicine. 33:282–288.

Nanchahal, J., C. Ball, I. Rombach, L. Williams, N. Kenealy, H. Dakin, H. O’Connor, D. Davidson, P. Werker, S.J. Dutton, M. Feldmann, and S.E. Lamb. 2022. Anti-tumour necrosis factor therapy for early-stage Dupuytren’s disease (RIDD): a phase 2b, randomised, double-blind, placebo-controlled trial. Lancet Rheumatol. 4:E407–e416.

Naylor, A.J., G. Desanti, A.N. Saghir, and R.S. Hardy. 2018. TNFα depleting therapy improves fertility and animal welfare in TNFα-driven transgenic models of polyarthritis when administered in their routine breeding. Lab Anim. 52:59–68.

Riester, S.M., D. Arsoy, E.T. Camilleri, A. Dudakovic, C.R. Paradise, J.M. Evans, J. Torres-Mora, M. Rizzo, P. Kloen, M.K. Julio, A.J. van Wijnen, and S. Kakar. 2015. RNA sequencing reveals a depletion of collagen targeting microRNAs in Dupuytren’s disease. BMC Med Genomics. 8:59.

Robbins, J.R., and K.G. Vogel. 1994. Regional expression of mRNA for proteoglycans and collagen in tendon. Eur J Cell Biol. 64:264–270.

Shih, B., S. Watson, and A. Bayat. 2012. Whole genome and global expression profiling of Dupuytren’s disease: systematic review of current findings and future perspectives. Ann Rheum Dis. 71:1440–1447.

Soreide, E., M.H. Murad, J.M. Denbeigh, E.A. Lewallen, A. Dudakovic, L. Nordsletten, A.J. van Wijnen, and S. Kakar. 2018. Treatment of Dupuytren’s contracture: a systematic review. Bone Joint J. 100-B:1138-1145.

Sykes, B., B. Puddle, M. Francis, and R. Smith. 1976. The estimation of two collagens from human dermis by interrupted gel electrophoresis. Biochem Biophys Res Commun. 72:1472–1480.

Takeda, K., A. Hatamochi, M. Arakawa, and H. Ueki. 1993. Effects of tumor necrosis factor-alpha on connective tissue metabolism in normal and scleroderma fibroblast cultures. Arch Dermatol Res. 284:440–444.

Tan, K., A.H.J. Withers, S.T. Tan, and T. Itinteang. 2018. The Role of Stem Cells in Dupuytren’s Disease: A Review. Plast Reconstr Surg Glob Open. 6:e1777.

Ten Dam, E.J., M.M. van Beuge, R.A. Bank, and P.M. Werker. 2016. Further evidence of the involvement of the Wnt signaling pathway in Dupuytren’s disease. J Cell Commun Signal. 10:33–40.

Tomasek, J.J., and C.J. Haaksma. 1991. Fibronectin filaments and actin microfilaments are organized into a fibronexus in Dupuytren’s diseased tissue. Anat Rec. 230:175–182.

van Beuge, M.M., E.J. Ten Dam, P.M. Werker, and R.A. Bank. 2016. Matrix and cell phenotype differences in Dupuytren’s disease. Fibrogenesis Tissue Repair. 9:9.

Vandesompele, J., K. De Preter, F. Pattyn, B. Poppe, N. Van Roy, A. De Paepe, and F. Speleman. 2002. Accurate normalization of real-time quantitative RT-PCR data by geometric averaging of multiple internal control genes. Genome Biology. 3:research0034.0031 - research0034.0011.

Varela-Rey, M., C. Montiel-Duarte, J.A. Osés-Prieto, M.J. López-Zabalza, J.P. Jaffrèzou, M. Rojkind, and M.J. Iraburu. 2002. p38 MAPK mediates the regulation of α1(I) procollagen mRNA levels by TNF-α and TGF-β in a cell line of rat hepatic stellate cells11The opinions or assertions contained herein are the private views of the authors and are not to be construed as official or as reflecting the views of the Department of the Army or the Department of Defense of the US. FEBS Letters. 528:133–138.

Verjee, L.S., K. Midwood, D. Davidson, D. Essex, A. Sandison, and J. Nanchahal. 2009. Myofibroblast distribution in Dupuytren’s cords: correlation with digital contracture. J Hand Surg Am. 34:1785–1794.

Verjee, L.S., J.S. Verhoekx, J.K. Chan, T. Krausgruber, V. Nicolaidou, D. Izadi, D. Davidson, M. Feldmann, K.S. Midwood, and J. Nanchahal. 2013. Unraveling the signaling pathways promoting fibrosis in Dupuytren’s disease reveals TNF as a therapeutic target. Proc Natl Acad Sci U S A. 110:E928–937.

Verrecchia, F., E.F. Wagner, and A. Mauviel. 2002. Distinct involvement of the Jun-N-terminal kinase and NF-kappaB pathways in the repression of the human COL1A2 gene by TNF-alpha. EMBO Rep. 3:1069–1074.

Vos, A.C., M.E. Wildenberg, M. Duijvestein, A.P. Verhaar, G.R. van den Brink, and D.W. Hommes. 2011. Anti-tumor necrosis factor-α antibodies induce regulatory macrophages in an Fc region-dependent manner. Gastroenterology. 140:221–230.

Worrell, M. 2012. Dupuytren’s disease. Orthopedics. 35:52–60.

Wynn, T.A. 2008. Cellular and molecular mechanisms of fibrosis. The Journal of pathology. 214:199–210.

Zeisberg, M., and R. Kalluri. 2013. Cellular mechanisms of tissue fibrosis. 1. Common and organ-specific mechanisms associated with tissue fibrosis. Am J Physiol Cell Physiol. 304:C216-225.

